# A mechanism to minimize errors during non-homologous end joining

**DOI:** 10.1101/563197

**Authors:** Benjamin M. Stinson, Andrew T. Moreno, Johannes C. Walter, Joseph J. Loparo

## Abstract

Enzymatic processing of DNA underlies all DNA repair, yet inappropriate DNA processing must be avoided. In vertebrates, double-strand breaks are repaired predominantly by non-homologous end-joining (NHEJ), which directly ligates DNA ends. NHEJ has the potential to be highly mutagenic because it employs DNA polymerases, nucleases, and other enzymes that modify incompatible DNA ends to allow their ligation. Using a biochemical system that recapitulates key features of cellular NHEJ, we show that end-processing requires formation of a “short-range synaptic complex” in which DNA ends are closely aligned in a ligation-competent state. Furthermore, single-molecule imaging directly demonstrates that processing occurs within the short-range complex. This confinement of end processing to a ligation-competent complex ensures that DNA ends undergo ligation as soon as they become compatible, thereby minimizing mutagenesis. Our results illustrate how the coordination of enzymatic catalysis with higher-order structural organization of substrate maximizes the fidelity of DNA repair.

## INTRODUCTION

Genome stability is central to the survival of all organisms. Faithful DNA replication and repair ensure genome stability, yet these processes employ scores of enzymes that chemically alter DNA and must be carefully regulated. A prominent example is non-homologous end joining (NHEJ), which repairs the majority of DNA double-strand breaks (DSBs) in vertebrate cells through direct re-ligation (Chiruvella et al., 2013; Karanam et al., 2012; Lieber, 2010; Radhakrishnan et al., 2014; Waters et al., 2014). DSBs arise from myriad spontaneous (e.g., reactive oxygen species, ionizing radiation) and programmed (e.g., V(D)J recombination, Cas9) sources, and failure to repair such DSBs results in cell death or malignancy-driving genomic rearrangements (Aparicio et al., 2014). Broken DNA ends are chemically diverse, and many are not compatible for immediate ligation. To solve this problem, NHEJ end-processing enzymes modify DNA ends until they are ligatable (Povirk, 2012). However, end processing also has the potential to introduce mutations by inserting or deleting nucleotides. Whereas this mutagenicity is exploited for immune receptor diversity and CRISPR-Cas9 gene disruption, it needs to be restrained during repair of unprogrammed breaks to prevent genome instability. At present, little is known about how DNA processing during NHEJ is regulated.

NHEJ is initiated by the ring-shaped Ku70/Ku80 heterodimer (Ku) (Mimori et al., 1986; Walker et al., 2001), which encircles broken DNA ends and recruits the DNA-dependent protein kinase catalytic subunit (DNA-PKcs) (Falck et al., 2005; Gottlieb and Jackson, 1993; Lees-Miller et al., 1990; Sibanda et al., 2017). DNA-PKcs phosphorylates multiple NHEJ factors, including itself, and its kinase activity is required for efficient end joining (Davis et al., 2014; Dobbs et al., 2010; Jette and Lees-Miller, 2015). Ends are ligated by a complex of DNA Ligase IV (Lig4) and XRCC4 (Grawunder et al., 1997), and joining is stimulated by the XRCC4 paralog XLF (Ahnesorg et al., 2006). Another XRCC4-like protein, PAXX (<u>Pa</u>ralog of <u>X</u>LF and <u>X</u>RCC4), appears to have functions overlapping with those of XLF (Kumar et al., 2016; Ochi et al., 2015; Tadi et al., 2016; Xing et al., 2015). These core factors cooperate with end-processing enzymes to allow joining of chemically incompatible ends: DNA polymerases (e.g., pol λ, pol μ) fill in overhangs and gaps (Bertocci et al., 2003, 2006; Lee et al., 2004; Ma et al., 2004; Mahajan et al., 2002); nucleases (e.g., Artemis) remove damaged nucleotides (Ma et al., 2002); polynucleotide kinase 3′-phosphatase (PNKP) adds 5′-phosphates and removes 3′ phosphates (Koch et al., 2004); tyrosyl-DNA phosphodiesterase 1 (Tdp1) removes topoisomerase I adducts (Yang et al., 1996) and other 3′ modifications (Inamdar et al., 2002; Interthal et al., 2005); tyrosyl-DNA phosphodiesterase 2 (Tdp2) removes 5′ DNA-topoisomerase II adducts (Gómez-Herreros et al., 2013; Ledesma et al., 2009); and aprataxin removes 5′ adenylate moieties resulting from abortive ligation (Ahel et al., 2006; Clements et al., 2004). Thus, processing enzymes equip NHEJ with numerous means to resolve end incompatibility.

NHEJ processing enzymes can introduce unnecessary mutations, leading to the widespread notion that NHEJ is highly error-prone and “sloppy.” For example, overexpression of site-specific endonucleases leads to the formation of NHEJ-dependent junctions with variable sequences (Lin et al., 1999; Lukacsovich et al., 1994; Phillips and Morgan, 1994; Rouet et al., 1994; Sargent et al., 1997). However, because accurate repair regenerates the endonuclease recognition sequence, multiple rounds of cutting and repair usually occur before an error arises. Consequently, the fidelity of repair is actually much higher than suggested by these studies. Indeed, when re-cleavage is disfavored or disallowed, compatible (i.e., blunt, complementary) ends are usually repaired without errors (Chiruvella et al., 2013; Karanam et al., 2012; Lieber, 2010; Radhakrishnan et al., 2014; Waters et al., 2014a). Moreover, processing of incompatible ends is limited to the minimum required for ligation (Bétermier et al., 2014; Budman and Chu, 2005; Guirouilh-Barbat et al., 2004; Waters et al., 2014b). Collectively, these studies suggest that NHEJ minimizes the extent of end processing and prioritizes ligation, but the molecular basis of this prioritization is unknown.

Coordination of end processing with synapsis, which aligns DNA ends for ligation, may play an important regulatory role. Using single-molecule Forster resonance energy transfer (smFRET) imaging, we previously showed that synapsis occurs in two stages (Figure 1A) (Graham et al., 2016): initially, Ku and DNA-PKcs form a “long-range” synaptic complex, in which dye-labeled DNA ends are tethered at a distance greater than that required for FRET (>100 Å); subsequently, DNA-PKcs kinase activity, XLF, and Lig4-XRCC4 convert the long-range complex into a high-FRET “short-range” complex in which ends are held close together in a ligation-competent state. Stable synapsis of DNA ends using purified human proteins shows similar requirements (Wang et al., 2018). Conflicting models have been proposed in which end processing occurs independently of synapsis (Lieber, 2010; Ma et al., 2004), while ends are synapsed (Conlin et al., 2017; Waters et al., 2014b), or upon dissolution of synapsis after failed ligation (Reid et al., 2017). However, without a means to simultaneously measure end synapsis and processing, distinguishing among these models has been impossible.

**Figure 1.**
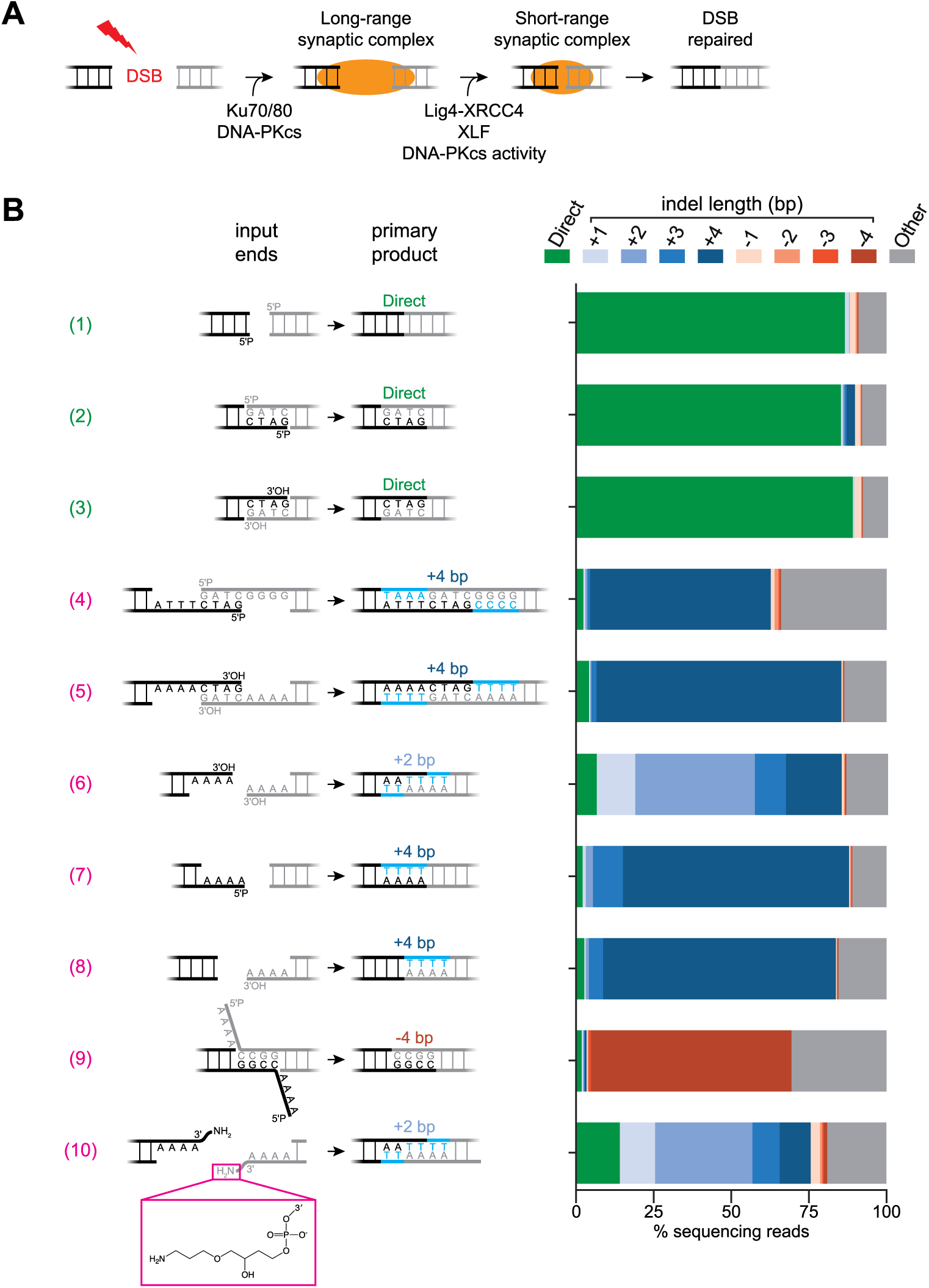
Joining of diverse end structures in *Xenopus* egg extracts. (A) Model of two-stage DNA end synapsis during NHEJ (Graham et al., 2016). (B) Linear DNA molecules with the indicated ends were added to egg extracts and incubated for 90 min. DNA was extracted, and joints were amplified by PCR and analyzed by paired-end Illumina sequencing. Indel length is relative to direct ligation of the top strand depicted for each end pair. Light blue sequence corresponds to nucleotides inserted during joining. Bar graphs were generated by identifying junction reads with the indicated indel length and normalizing to the total number of reads. See Methods for controls regarding fidelity of amplification, library preparation, and sequencing. Analysis of aligned sequencing reads is reported in Data S1.

Here, we use single-molecule fluorescence imaging in *Xenopus* egg extracts to observe DNA end synapsis and processing in real time. We first establish that egg extracts support ligation of chemically diverse DNA ends, and that joining of each type of incompatible DNA end requires a distinct processing enzyme. Next, we show that the activities of all processing enzymes examined require Lig4-XRCC4, XLF, and DNA-PKcs kinase activity, establishing a dependence of processing on formation of the short-range synaptic complex. Moreover, smFRET assays that simultaneously monitor synapsis and end-processing demonstrate that processing occurs preferentially in the short-range synaptic complex. Our results establish a model in which confining end processing to the Lig4-dependent short-range synaptic complex promotes immediate ligation of compatible ends and ensures that incompatible ends are ligated as soon as they become compatible, thereby minimizing end processing.

## RESULTS

### Egg extracts recapitulate diverse end-processing events

To investigate the regulation of processing, we used *X. laevis* egg extracts, which join DNA ends in a manner dependent on Ku, DNA-PKcs, Lig4-XRCC4, and XLF (Graham et al., 2016; Labhart, 1999; Pfeiffer and Vielmetter, 1988; Thode et al., 1990; Virgilio and Gautier, 2005). We generated a series of linear DNA substrates with compatible or incompatible ends, added these substrates to egg extracts, and sequenced the resulting NHEJ repair junctions (Figures 1B and S1A). We verified that a representative product spectrum was maintained through amplification and sample processing (see Methods).

Compatible ends (i.e., blunt, complementary) were typically ligated without processing (Figure 1B, #1-3), underscoring that NHEJ is not intrinsically mutagenic. Due to minor contaminants in the input DNA and errors introduced during sample preparation, these measurements represent a lower limit of NHEJ fidelity (see Methods). Remarkably, incompatible ends underwent a variety of processing events prior to joining, including gap-filling (Figure 1B, #4-8), nucleolytic trimming (Figure 1B, #9), and 3′-adduct removal (Figure 1B, #10). The extent of processing was generally limited to a few nucleotides, and most substrates were joined with a single predominant sequence, even in the absence of microhomology to guide end alignment.

To identify end-processing enzymes in this cell-free system, we added linear DNA substrates containing radioactively labeled DNA ends (Figure S1A) to egg extract that had been immunodepleted of specific factors. At different time points, the DNA was recovered, digested with SacI and KpnI, separated by denaturing PAGE, and the starting material, processing intermediates, and ligation products were visualized at nucleotide resolution by autoradiography (Figure 2A). Immunodepletion of pol λ and pol μ prevented gap-filling of DNA ends with paired primer termini (Figures 2A, lanes 1-8, and S1B). Re-addition of recombinant pol λ but not pol μ restored efficient gap-filling of these DNA ends (Figure 2A, lanes 9-16), consistent with previous reports (McElhinny et al., 2005). Immunodepletion of pol λ and pol μ also prevented gap-filling of DNA ends with unpaired primer termini (Figure S1C, lanes 1-8); conversely, re-addition of pol μ but not pol λ supported gap-filling of these ends (Figure S1C, lanes 9-16). Tdp1 efficiently removed a 3′-phosphodiester adduct (Figure 2B and S1D; see Figure 1B, #10 for adduct structure), consistent with its broad substrate specificity (Interthal et al., 2005). PNKP was essential for phosphorylation of 5′-hydroxyl ends prior to joining (Figures 2C and S1E). Although 5′ flaps were removed prior to joining (Figure 1B, #9), immunodepletion of Artemis, a nuclease implicated in NHEJ (Chang et al., 2015), had no effect on joining of these ends (Figure S1F), suggesting involvement of an unidentified nuclease. PNKP, pol λ, pol μ, and Tdp1 were not required for joining of compatible ends, indicating that these enzymes are dispensable for the core NHEJ reaction (Figure S1G). Thus, as seen in human cells (Guirouilh-Barbat et al., 2004; Waters et al., 2014b), *X. laevis* egg extracts support efficient end processing by diverse enzymes but employ this mechanism only when necessary.

**Figure 2.**
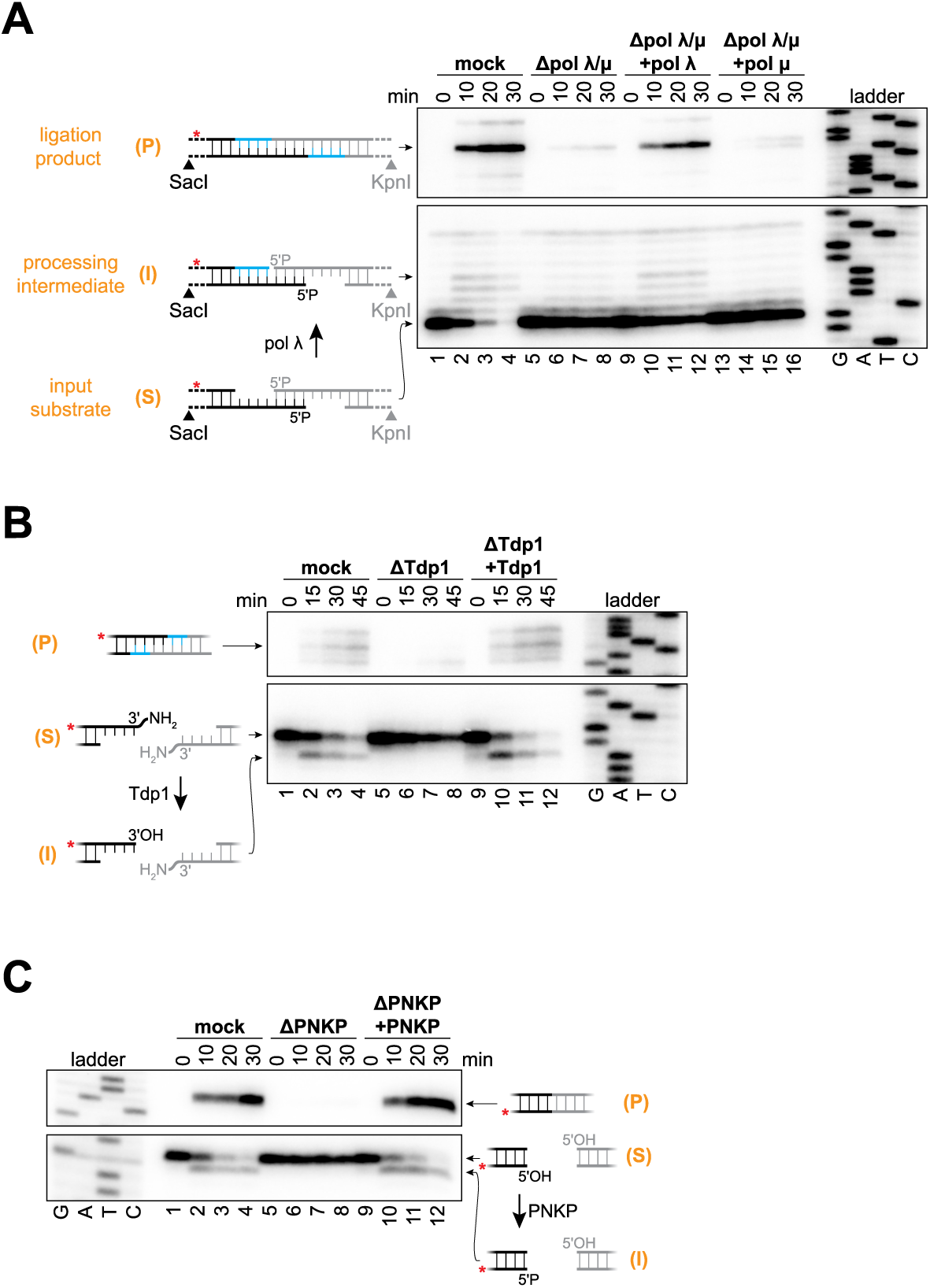
Involvement of pol λ and Tdp1 in end processing. (A to C) Radiolabeled linear DNA molecules with the depicted ends were added to the indicated egg extracts, and reaction samples were stopped at the indicated time points. DNA was extracted, digested, and analyzed by denaturing PAGE and autoradiography. Lower panels, un-joined DNA ends; upper panels, joined DNA ends. Red asterisk indicates radiolabeled strand. Immunodepletion of pol λ alone blocked joining of the ends depicted in (A) (see Figure S1G, lanes 22 to 24). Immunodepleted extracts were supplemented with the following final concentrations of recombinant protein as indicated: pol λ and pol μ, 10 nM; Tdp1, 40 nM. S, input substrate; I, processing intermediate; P, ligation product.

### Blocking short-range synapsis inhibits end processing

We reasoned that end processing might be regulated through coordination with synapsis. Based on our two-stage model (Figure 1A) (Graham et al., 2016), we hypothesized that end processing occurs only in the Lig4-dependent short-range synaptic complex. Such a mechanism would promote immediate ligation of compatible ends and ensure that incompatible ends undergo ligation as soon they become compatible, thereby minimizing processing.

A prediction of this hypothesis is that disruptions of short-range synapsis should block end processing. To test this idea, we used three independent means—Lig4-XRCC4 immunodepletion, XLF immunodepletion, or DNA-PKcs inhibition—to block short-range synapsis and asked whether end processing was inhibited. Single-molecule experiments (see below) verified that each of these perturbations blocked short-range synapsis of the pol λ substrate ends shown in Figure 2A (Figure S2A), as reported previously for blunt ends (Graham et al., 2016). Immunodepletion of XRCC4 (Figure S2B), which also depletes Lig4 (Graham et al., 2016), inhibited gap filling by pol λ, as well as subsequent ligation (Figure 3A, lanes 1 to 8). Gap filling was restored by re-addition of recombinant wild-type Lig4-XRCC4 (Figure 3A, lanes 9 to 12). Re-addition of catalytically-inactive Lig4-XRCC4, which promotes short-range synapsis (Graham et al., 2016), also restored gap filling, resulting in accumulation of a processed but unligated intermediate (Figure 3A, lanes 13 to 16). Catalytically-inactive Lig4-XRCC4 supported a low level of joining (Figure 3A, compare lanes 13 to 16 with lanes 5 to 8), likely because it promotes generation of compatible ends that can be joined by residual endogenous Lig4-XRCC4 or another ligase. Like Lig4-XRCC4, XLF and DNA-PKcs kinase activity were also required for gap filling (Figures 3A, lanes 17 to 32, and S2B). We next asked whether short-range synapsis is generally required for processing. Strikingly, 3′-adduct removal by Tdp1 (Figure 3B), gap-filling by pol μ (Figure S2C), 5′-hydroxyl phosphorylation by PNKP (Figure S2D), and nucleolytic flap removal (Figure S2E) all depended on Lig4-XRCC4, XLF, and DNA-PKcs kinase activity. Thus, three independent perturbations of short-range synapsis substantially inhibited every processing activity measured, strongly suggesting that end processing generally depends on short-range synapsis.

**Figure 3.**
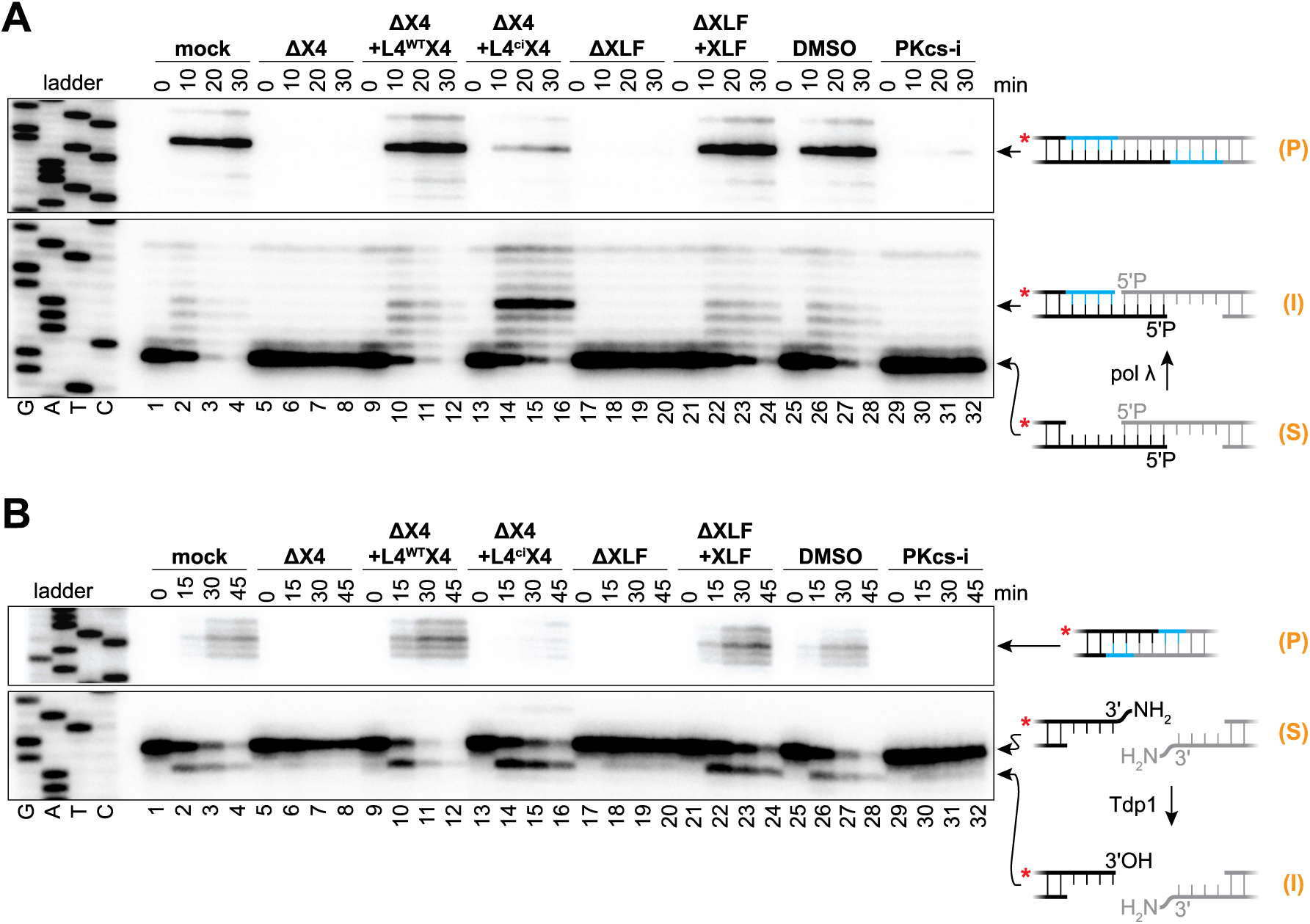
Disruption of short-range synapsis inhibits end processing by pol λ and Tdp1. (A and B) Denaturing PAGE analysis of processing and joining of depicted DNA ends in indicated extracts as in Figure 2. X4, XRCC4; L4^WT^ and L4^ci^, wild-type and catalytically-inactive Lig4, respectively. Recombinant Lig4-XRCC4 and XLF were added to final concentrations of 50 nM and 100 nM, respectively. PKcs-i in DMSO was added to a final concentration of 50 μM. S, input substrate; I, processing intermediate; P, ligation product.

### Single-molecule imaging of NHEJ polymerase activity

Our results are consistent with the idea that end processing occurs within the short-range complex. However, they do not exclude a model (Reid et al., 2017) in which incompatible DNA ends dissociate from the short-range complex before undergoing processing, followed by subsequent synapsis and ligation attempts. To distinguish between these models, we performed single-molecule FRET analysis to simultaneously monitor DNA-end synapsis and processing in real time. We attached a ~3 kb linear DNA molecule with partially-complementary 5′ overhangs to a passivated glass coverslip in a microfluidic channel (Figure 4A). To monitor short-range synapsis using FRET, we placed a donor fluorophore near one DNA end and an acceptor fluorophore near the other end (Graham et al., 2016), which did not affect joining (Figure S3A). The extract was desalted and supplemented with dUTP labeled with the dark quencher BHQ-10 (dUTP^Q^; Figure S3B) so that dUTP^Q^ incorporation opposite a templating adenosine would quench donor fluorescence (Figure 4A). dATP, dGTP, dCTP, and ATP were also added to the desalted extract to support gap filling and joining. Ensemble experiments demonstrated that dUTP^Q^ was efficiently incorporated during joining of these ends (Figure S3C, lanes 5 to 8). Figure 4B shows a representative single-molecule fluorescence trajectory of joining in the presence of dUTP^Q^. After addition of extract, we observed a transition from a low-FRET state to a high-FRET state, indicating short-range synapsis had occurred, followed by quenching in the high-FRET state, consistent with dUTP^Q^ incorporation. Subsequent fluorescence fluctuations allowed us to unambiguously identify bona fide quenching events and exclude donor photobleaching. We ascribe the periodic return of donor fluorescence to quencher “blinking,” rather than repeated futile dUTP^Q^ incorporation attempts, as a minimal reconstitution showed fluorescence fluctuations following the initial quenching event even after polymerase was removed (Figure S3D-S3L).

**Figure 4.**
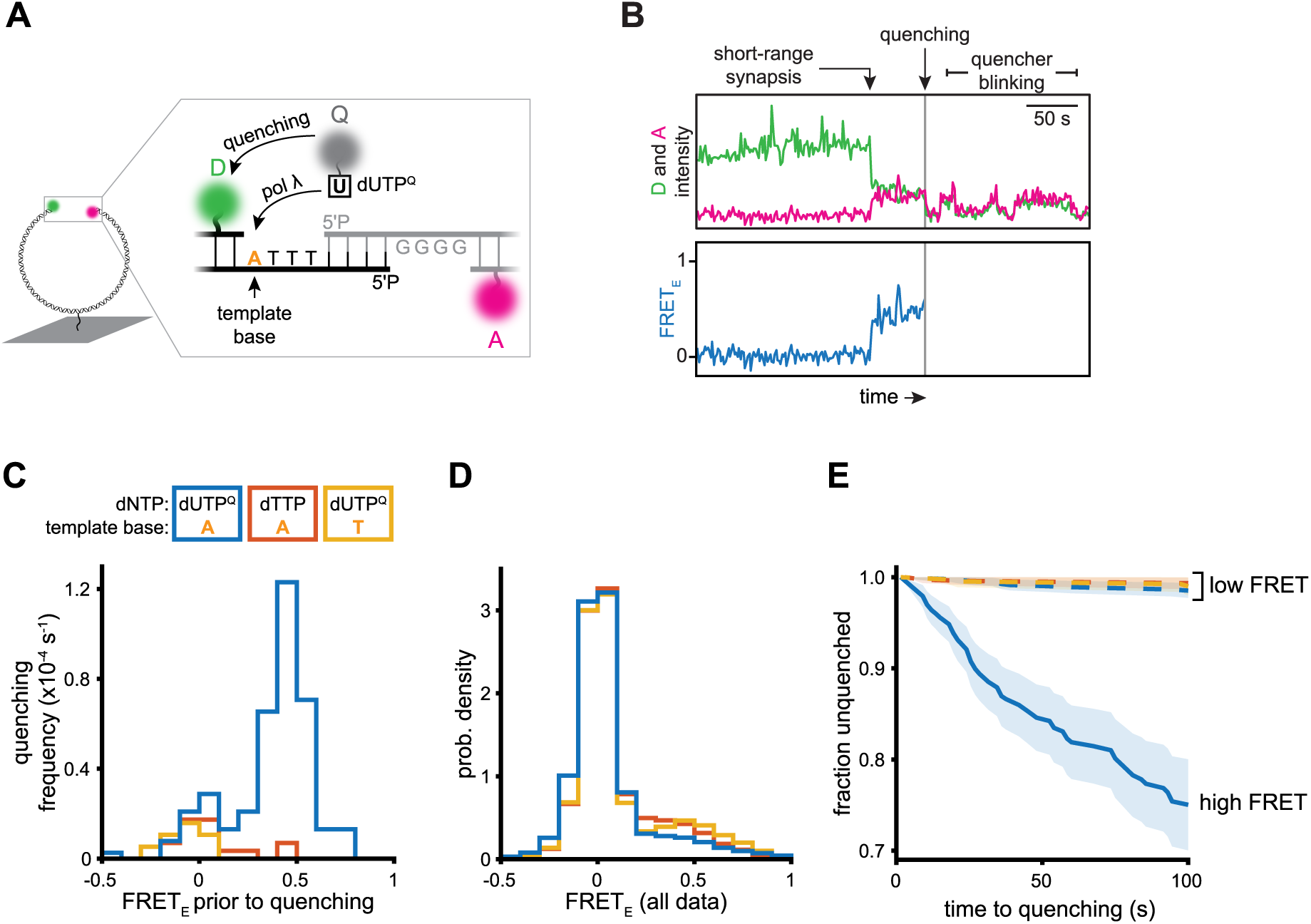
pol λ acts within the short-range synaptic complex. (A) Experimental scheme for single-molecule assay of DNA-end synapsis and polymerase activity. D, Cy3 donor fluorophore; A, Cy5 acceptor fluorophore; Q, BHQ10 quencher. Fluorophore positions are noted in Table S1. (B) Example trajectory. The Cy3 donor fluorophore was excited and donor emission (green) and Cy5 acceptor emission (magenta) were recorded (upper panel) and used to calculate FRET efficiency (blue, lower panel). (C) Frequency of quenching events as a function of FRET efficiency immediately prior to quenching under the indicated conditions. Blue, extracts supplemented with dUTP^Q^, adenine DNA template base (see (A)) substrate (*n* = 145 events from 9 independent experiments); red, extracts supplemented with dTTP, adenine DNA template base substrate (*n* = 16 events from 6 independent experiments); yellow, extracts supplemented with dUTP^Q^, thymine DNA template base (*n* = 8 events from 6 independent experiments). See Methods for calculation of quenching frequency and Figure S4A for kinetic analysis and histograms represented as probability density. (D) Histogram of the FRET efficiency of all frames for all molecules prior to censoring. Colors as in C. (E) Quenching survival kinetics (Kaplan-Meier estimate) under the indicated conditions (colors as in C). The x-axis indicates dwell time in the high-FRET (solid line) or low-FRET (dashed lines) state prior to quenching. Shaded areas, 95% confidence intervals.

Importantly, the vast majority (~80%) of quenching events occurred in the high-FRET state (FRET_E_ ≈ 0.5; Figure 4C, blue) even though the low-FRET state was much more abundant overall (FRET_E_ ≈ 0; Figure 4D, blue). Inclusion of dTTP instead of dUTP^Q^ revealed a background frequency of spuriously-detected low-FRET quenching events (Figure 4C, red) similar to that observed when dUTP^Q^ was present (Figure 4C, blue). Thus, dUTP^Q^ was efficiently incorporated in the high-FRET state, and the rate of incorporation in the low-FRET state was indistinguishable from the background rate (Figure 4E). Changing the template base from adenine to thymine, inclusion of dTTP instead of dUTP^Q^, combined immunodepletion of pol λ and pol μ, or inhibition of DNA-PKcs all suppressed dUTP^Q^ incorporation (Figure S3C, lanes 9 to 28) and high-FRET quenching events (Figures 4C, yellow,S4B and S4C). We conclude that pol λ activity is tightly coupled to the short-range synaptic complex.

### Single-molecule imaging of Tdp1 activity

We next asked whether Tdp1 also acts in the short-range synaptic complex. To measure Tdp1 activity on single DNA molecules, we conjugated the 3′ adduct removed by Tdp1 (Figure 2B) to a donor fluorophore (Figure 5A). A DNA substrate with this modification on one end and an acceptor fluorophore in the duplex region of the other end allowed observation of short-range synapsis via FRET and Tdp1 activity via loss of donor signal (Figure 5A). Ensemble experiments verified that the fluorescent 3′ adduct was efficiently removed prior to joining (Figure S5A, lanes 1 to 4). To increase the efficiency of NHEJ-mediated synapsis in single-molecule experiments, we blocked the competing resection pathway by depleting extracts of the Mre11 nuclease, which did not significantly affect end processing or joining (Figure S5A, lanes 5 to 8). We frequently observed a transition from a low-FRET state to a high-FRET state followed by a loss of donor signal (e.g., Figure 5B). The majority (~60%) of donor loss events occurred in the high-FRET state (Figure 5C, blue) despite its overall low abundance (~20% of all frames; Figure 5D, blue). Thus, donor loss occurred ~5-fold more quickly in the high-FRET state compared to the low-FRET state (Figure 5E, solid blue vs. dashed blue). Tdp1 immunodepletion greatly reduced the frequency of donor loss events from both low- and high-FRET states (Figure 5C, red), and this reduction was rescued upon re-addition of recombinant Tdp1 (Figure 5C, yellow), indicating that Tdp1 and not photobleaching was responsible for the vast majority of detected donor loss events. The occasional donor loss events observed in the low-FRET state likely involve Tdp1 acting on transiently unprotected ends (see below). In summary, all processing reactions measured depended on formation of the short-range complex (Figures 3 and S2), and in at least two of these—gap filling and adduct removal—end processing occurred preferentially in the high FRET state (Figures 4 and 5). Together, the data suggest that processing generally occurs in the short-range complex.

**Figure 5.**
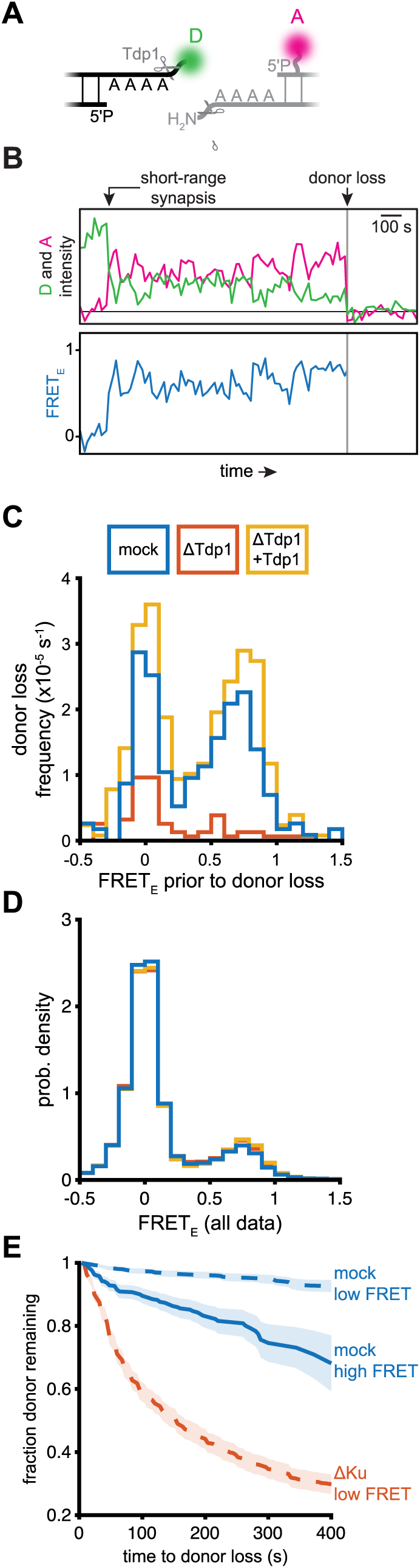
Short-range synapsis accelerates Tdp1 activity. (A) Experimental scheme for single-molecule assay of Tdp1 activity and DNA-end synapsis. The DNA substrate was a derivative of the 3′-adduct substrate depicted in Figure 2B (see Figure 1, #10 for chemical structure), with Cy3B conjugated to one of the 3′-adducts. D, Cy3B donor fluorophore; A, ATTO647N acceptor fluorophore. (B) Example trajectory. The Cy3B donor fluorophore was excited and donor emission (green) and ATTO647N acceptor emission (magenta) were recorded (upper panel) and used to calculate FRET efficiency (blue, lower panel). (C) Frequency of donor loss events as a function of FRET efficiency immediately prior to donor loss under the indicated conditions. Blue, mock-depleted extracts (*n* = 222 events from 2 independent experiments); red, Tdp1-depleted extracts (*n* = 65 events from 2 independent experiments); yellow, Tdp1-depleted extracts supplemented with 40 nM recombinant Tdp1 (*n* = 339 events from 2 independent experiments). See Methods for calculation of donor loss frequency and Figure S5D-S5F for kinetic analysis and histograms represented as probability density. (D) Histograms of FRET efficiency of all frames for all molecules prior to censoring. Colors as in C. (E) Donor survival kinetics (Kaplan-Meier estimate) under the indicated conditions. The x-axis indicates dwell time in the high-FRET (solid line) or low-FRET (dashed lines) state prior to donor loss. Blue, donor survival kinetics in mock-depleted extract (*n* = 76 events, dashed line; 107 events, solid line); red, donor survival kinetics in Ku-depleted extract (*n* = 687 events); shaded areas, 95% confidence intervals.

### Ku protects ends from bound processing factors and off-pathway enzymes

We next addressed how end processing is prevented before formation of the short-range synaptic complex. One possibility is that short-range synapsis is required to recruit end-processing enzymes. To test this idea, we attached DNA to magnetic beads and isolated bound end-processing factors in the presence or absence of DNA-PKcs inhibitor, which blocks short-range synapsis (Figures 6A and S6A). However, as shown in Figure 6A, PNKP, pol λ, pol μ, and Tdp1 were each enriched on DSB-containing DNA relative to the intact control whether or not the short-range synaptic complex was allowed to form, suggesting that end-processing enzymes are recruited prior to short-range synapsis.

**Figure 6.**
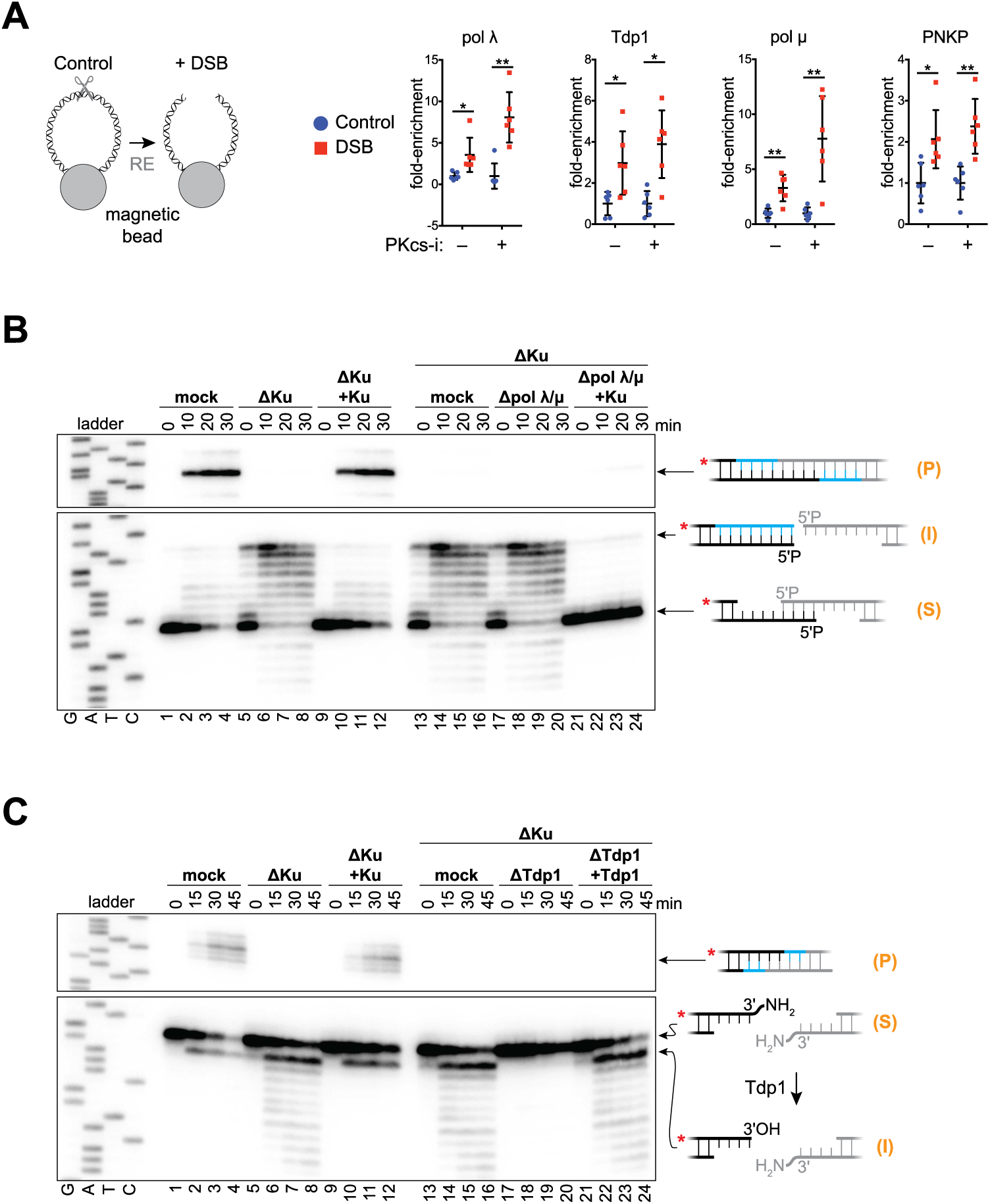
Ku protects DNA ends from premature and off-pathway processing. (A) Western blot analysis of end-processing enzyme recruitment to a DSB (blunt ends introduced by XmnI digestion) in mock-treated or DNA-PKcs-inhibited extracts (see Figure S6 for a representative experiment and explanation of quantification). Fold-enrichment is relative to undigested DNA (*n* = 6, mean ± s.d., \**p* < 0.001, \*\**p* < 0.001, two-tailed *t*-test). (B and C) Denaturing PAGE analysis of end processing and joining of the depicted DNA ends in the indicated extracts as in Figure 2. Recombinant Ku and Tdp1 were added as indicated to final concentrations of 300 nM and 40 nM, respectively. S, input substrate; I, processing intermediate; P, ligation product.

Based on the above results, we postulated that prior to short-range synapsis, core NHEJ factors protect DNA ends from the bound processing factors. Indeed, immunodepletion of Ku (Figure S6B) caused aberrant fill-in of 5′ overhang ends without subsequent joining (Figure 6B, lanes 5 to 8), consistent with a role for Ku in DNA end protection (Liang and Jasin, 1996). Re-addition of recombinant Ku restored typical gap-filling and joining (Figure 6B, lanes 9 to 12). Importantly, aberrant fill-in occurred even when pol λ and pol μ were immunodepleted in combination with Ku, suggesting a non-NHEJ polymerase can fill in 5′ overhangs when Ku is absent (Figures 6B, lanes 17 to 20, and S6C). Addition of recombinant Ku to the Ku/pol λ/pol μ-depleted extract did not rescue gap-filling and joining (Figures 6B, lanes 21-24), indicating that pol λ, which is required for NHEJ of these ends (Figure 2A), was functionally depleted in this experiment. Ku also suppressed aberrant 3′→5′ exonuclease activity following 3′ adduct removal (Figure 6C, lanes 5 to 12). Combined immunodepletion of Ku and Tdp1 (Figure S6D) prevented both 3′-adduct removal and exonuclease activity (Figure 6C, lanes 17 to 20), and re-addition of recombinant Tdp1 restored these activities (Figure 6C, lanes 21 to 24). Thus, Tdp1 can access DNA ends in the absence of Ku, but ends are protected in the presence of Ku when short-range synapsis is blocked (Figure 3B). Additionally, Ku depletion resulted in aberrant nuclease activity on blunt ends, 3′-overhang ends lacking microhomology, and 5′-overhang ends with internal microhomology (Figure S6E-S6G). We conclude that, prior to short-range synapsis, Ku protects DNA ends from processing by both NHEJ and off-pathway enzymes, and ends become accessible for processing upon short-range synapsis.

As shown above, Tdp1 had measurable activity outside of the short-range synaptic complex (Figures 5C and S5B). We postulated that Tdp1 acts on DNA ends if they are transiently unprotected prior to short-range synapsis. Consistent with this idea and the above ensemble experiments (Figure 6C), Ku immunodepletion in the single-molecule Tdp1 assay (Figure 5A) resulted in rapid donor loss from the low-FRET state (Figures 5E, red dashed line, and S5C, red), and this effect was suppressed by re-addition of recombinant Ku (Figure S5C, yellow). Tdp1 has known roles outside of NHEJ and does not typically alter DNA sequence (El-Khamisy and Caldecott, 2006). These traits may explain the less stringent coupling of Tdp1 activity to the formation of the short-range synaptic complex.

## DISCUSSION

Our findings reveal how NHEJ minimizes mutagenesis without sacrificing the versatility needed to repair diverse DSBs (Figure 7). Both before ends undergo synapsis and after formation of the long-range synaptic complex, Ku protects DNA ends from unrestricted processing. Thereafter, formation of the short-range synaptic complex potentiates the action of the DNA end processing machinery. Confining processing events to the short-range complex minimizes mutagenesis in multiple ways. First, because Lig4 is required for short-range synapsis, compatible DNA ends have a high probability to undergo ligation without processing, as seen in extracts (Figure 1, #1-3) and in cells (Waters et al., 2014b). Second, as soon as incompatible ends become compatible, they undergo ligation, avoiding unnecessary processing. For example, in Figure 1B #9, nuclease activity is limited to flap removal. Third, short-range synapsis promotes utilization of pre-existing microhomology (Figure 1B, #4, #5, and #9). This feature helps preserve the original sequence in cases where DSBs retain base complementarity but have damaged, unligatable termini, such as 3′-phosphoglycolates introduced by ionizing radiation (Povirk, 2012). Finally, maintenance of short-range synapsis during processing avoids unpaired-end intermediates that may give rise to translocations. Notably, in human cell extracts, Lig4-XRCC4, XLF, and/or DNA-PKcs activity have been implicated in NHEJ polymerase and nuclease activity (Akopiants et al., 2009; Budman et al., 2007; Lee et al., 2003, 2004), consistent with a similar regulation of end processing in human cells.

**Figure 7.**
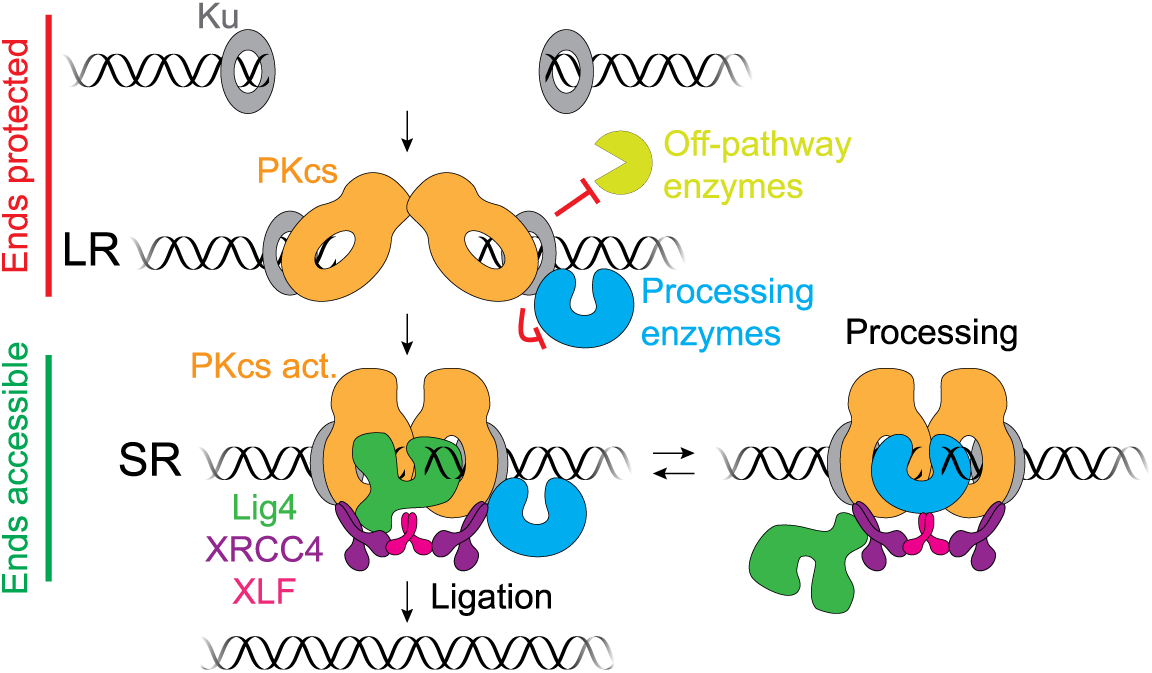
Model of DNA end processing during NHEJ.

### Lig4-XRCC4 dynamics during joining of incompatible ends

To explain how ligation is prioritized over end processing, we propose that Lig4 directly engages DNA ends to drive short-range synapsis (Figure 7). Such DNA bridging may constitute the non-catalytic role of Lig4 in synapsis that we and others have observed (Cottarel et al., 2013; Graham et al., 2016). In this way, ends are protected by Ku prior to short-range synapsis, and Lig4 itself acts as an end protection factor in the short-range synaptic complex by excluding processing enzymes before ligation has been attempted. To facilitate processing of incompatible ends, we propose that Lig4 dynamically releases DNA ends to allow processing and promptly re-engages them to ensure ligation of minimally processed ends (Figure 7). We recently reported that a single XLF dimer interacts with two XRCC4-Lig4 complexes to mediate short-range synapsis (Figure 7) (Graham et al., 2018). Thus, after initial DNA end binding by Lig4, an XRCC4-XLF-XRCC4 “bridge” may maintain short-range synapsis, allowing Lig4 to release DNA ends but remain within the repair complex via its XRCC4 interaction. Alternatively, short-range synapsis may be established and maintained by another non-catalytic function of Lig4, such as XRCC4 recruitment. A recent study suggested that Lig4 must maintain direct interactions with DNA ends during end processing in a manner that requires a Lig4-specific structural motif, insert1 (Conlin et al., 2017). However, it is unclear how Lig4 would be able to both maintain stable interactions with DNA ends and simultaneously allow productive binding of processing factors. Instead, we propose that insert1 enables Lig4 to bridge incompatible DNA ends for initial short-range synapsis, and that Lig4 then releases DNA ends to allow processing as described above. End protection by Ku and probably DNA-PKcs must also be relieved to allow for processing and ligation. This deprotection may occur though Ku translocation away from the break (Yoo and Dynan, 1999) and/or autophosphrylation-induced conformational changes or dissociation of DNA-PKcs (Chan and Lees-Miller, 1996; Cui et al., 2005; Ding et al., 2003; Dobbs et al., 2010; Goodarzi et al., 2006; Reddy et al., 2004). In summary, our results imply that mutagenesis during NHEJ is minimized by carefully restricting the access of processing factors to DNA ends, both before and after the ends have undergone synapsis.

### Hierarchical access of end-modifying enzymes

Our results provide two lines of evidence that NHEJ minimizes errors not only by prioritizing ligation, but also by utilizing processing enzymes hierarchically, as recently proposed (Strande et al., 2012; Waters et al., 2014a). First, we find that conservative end-processing events are favored over sequence-altering events. For example, ends requiring only 5′ phosphorylation for ligation are almost always joined without insertions or deletions (Figure 2C, lanes 1-4). Moreover, when PNKP is absent, such ends are not subject to significant polymerase or nuclease activity (Figure 2C, lanes 5-8). Thus, PNKP sits atop the processing hierarchy for these ends. Similarly, Tdp1 activity is preferred to nuclease activity in the removal of 3′ adducts (Figure 2B). Second, among sequence-altering end-processing events, polymerase fill-in was favored over nuclease activity. In principle, non-complementary ends could be rendered ligatable either through polymerase or nuclease activity, yet we observed a strong preference for polymerase activity (Figure 1B, #4-9). Together, these features may contribute to repair fidelity as depicted in Figure S7A. Lig4, PNKP, and Tdp1 repair ends without altering sequence and are positioned at the top of the utilization hierarchy. In the next tier, pol λ and pol μ alter sequence, but are limited in the extent of modification by the availability of single-stranded template (pol μ has reported template-independent synthesis activity, but it is much less efficient than templated synthesis (McElhinny et al., 2005)). Finally, nucleases could theoretically resolve any form of damage found at a DNA end, but they are utilized as a last resort owing to their mutagenicity. Future experiments will be necessary to determine the mechanism underlying this hierarchy. Together, confinement of end processing to the short-range synaptic complex and hierarchical utilization of processing enzymes finely tune processing of DNA ends to minimize errors during DSB repair via NHEJ.

### Regulation of end processing in V(D)J recombination

V(D)J recombination assembles immune receptor genes by introducing programmed DSBs that are repaired by NHEJ (Figure S7B) (Alt et al., 2013; Gellert, 2002). “Coding ends” generated by the RAG recombinase are joined to form the exon encoding the antigen-binding region. Coding ends exhibit substantial variation in the number of nucleotides inserted or deleted prior to joining (Figure S7C), thereby augmenting immune receptor diversity. How NHEJ promotes highly mutagenic processing of coding ends but not of spontaneous DSBs is an important question. Because the same core NHEJ factors direct both processes, we hypothesize that the overarching regulation of end processing is similar, but that V(D)J recombination incorporates unique DNA end structures and processing enzymes to promote junction variability. For example, RAG generates a hairpin terminus at coding ends, which the Artemis nuclease must cut before ligation can be attempted. Thus, whereas ligation is usually attempted before end processing for spontaneous DSBs, coding ends always require initial processing. Moreover, Artemis preferentially cleaves the hairpin two nucleotides 3′ of the hairpin tip, generating four-nucleotide 3′ overhangs that are usually non-complementary (Ma et al., 2002; Schlissel, 1998). Of the ends tested in this study, such a substrate had the most variable junctions (Figure 1B, #6), although this variability was still largely limited to 1-4 nucleotide insertions mediated by pol μ. In addition, V(D)J recombination utilizes the lymphocyte-specific terminal deoxynucleotidyl transferase (TdT), which has robust template-independent synthesis activity on 3′ overhangs (Deng and Wu, 1983). Interestingly, signal ends (Figure S7C), which are blunt and therefore do not require Artemis activity and do not present the preferred TdT end structure, are typically joined without any insertions or deletions (Gellert, 2002), similar to compatible spontaneous DSBs (Figure 1B, #1-3). Alternatively, coding end processing could be regulated by an entirely distinct mechanism. For example, based on reconstitution with purified NHEJ factors, it has been proposed that each DNA end independently recruits processing factors stochastically until ends become compatible for ligation (Lieber, 2010; Ma et al., 2004). Ultimately, to resolve how end processing during V(D)J recombination is uniquely regulated to generate junctional diversity, it will be critical to understand how RAG transfers coding ends to the NHEJ machinery (Bertocci et al., 2003, 2006; Lee et al., 2004b; Ma et al., 2004; Mahajan et al., 2002).

### Gene editing

Because NHEJ is the predominant pathway for repair of CRISPR-Cas9 DSBs, our results also have important implications for genome engineering. Efforts to improve the efficiency of CRISPR-Cas9-mediated gene disruption by overexpressing DSB processing enzymes (Chari et al., 2015) have been only modestly effective, most likely because such an approach does not circumvent DNA end protection before short-range synapsis or immediate ligation of compatible ends upon short-range synapsis. Our model predicts that a more effective approach will be to increase DNA end accessibility prior to ligation, thereby facilitating activity of processing enzymes.

### Conclusion

In summary, we find that the integration of DNA processing with formation of higher-order DNA repair complexes helps to insure the fidelity of NHEJ. A similar strategy might govern other crucial events in DNA repair, such as 5′→3′ resection of DSBs, which initiates homologous recombination (HR) and alternative end joining (Symington and Gautier, 2011). For example, tethering two broken DNA ends could trigger their simultaneous resection (Andres et al., 2015; Westmoreland and Resnick, 2013). In addition, resection may depend on tethering a broken end to its intact sister chromatid (Williams et al., 2008; Zhu et al., 2018), preventing excessive resection when a homologous repair template is not available. For these and other questions in the regulation of DNA metabolism, approaches that correlate structural features of protein-DNA complexes with DNA modification will serve as powerful tools.

---

**Figure S1.**
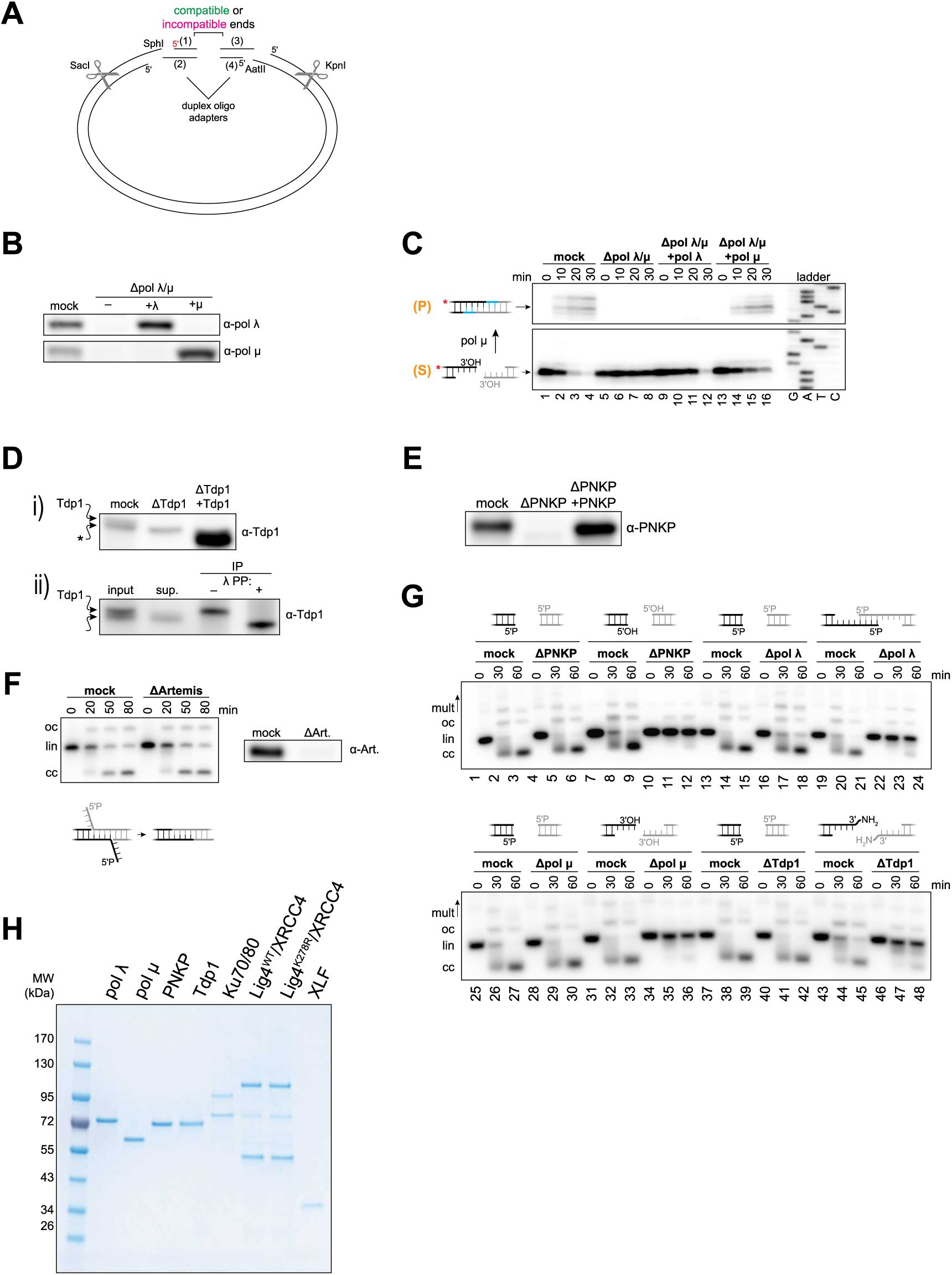
Identification of end-processing enzymes in *Xenopus* egg extracts. Related to Figures 1 and 2. (A) Schematic for generation of linear DNA molecules with variable end structures, as described in detail in the Methods. Four short synthetic oligonucleotides were used to generate two duplex adapters. One adapter contained SphI-compatible overhangs and the other contained AatII-compatible overhangs. The adapters were ligated to a 3-kb plasmid previously digested with SphI and AatII. Radiolabeled DNA substrates were typically generating by incorporating a ^32^P 5′ phosphate on one oligo (red). After incubation in extract, DNA was isolated and digested with SacI and KpnI before denaturing PAGE and autoradiography. (B) Western blot analysis of immunodepletions in Figures 2A and S1C. (C) Denaturing PAGE analysis of processing and joining of depicted DNA ends in indicated extracts as in Figure 2A. Recombinant pol λ and pol μ were added to a final concentration of 10 nM as indicated. Immunodepletion of pol μ alone blocked joining of the ends depicted in A and B, respectively (see Figure S1G, lanes 34 to 36). S, input substrate; P, ligation product. (D) Western blot analysis of immunodepletions in Figure 2B. (i) the Tdp1 antibody recognized a nonspecific band (asterisk) that migrated slightly faster and was poorly resolved from endogenous Tdp1. Recombinant Tdp1 migrated slightly faster than endogenous Tdp1. (ii) Endogenous Tdp1 was immunoprecipitated from extract and treated with λ protein phosphatase (λ PP), resulting in faster Tdp1 migration similar to the recombinant protein. Thus, lack of phosphorylation of the recombinant protein may necessitate the higher concentration required for robust Tdp1 activity. Sup., supernatant; IP, immunoprecipitate. (E) Denaturing PAGE analysis of processing and joining of depicted DNA ends in indicated extracts as in Figure 2. Bottom panel, Western blot analysis of immunodepletions. Recombinant PNKP was added to a final concentration of 50 nM as indicated. (F) A radiolabeled linear DNA molecule bearing the depicted 5′-flap ends was added to mock- or Artemis depleted extract. Reactions samples were stopped at the indicated time points and analyzed by native agarose gel electrophoresis and autoradiography. mult, multimers; oc, open circular plasmid; lin, linear; cc, closed circular plasmid. Right panel, Western blot analysis of immunodepletions. oc, open circular plasmid; lin, linear; cc, closed circular plasmid. (G) Extracts were immunodepleted of the indicated end-processing factors and tested for joining of blunt, phosphorylated ends that do not require processing. As a positive control, the same extracts were also tested for joining of ends requiring processing by the immunodepleted enzyme. Radiolabeled linear DNA molecules with the depicted ends were added to the indicated egg extracts and analyzed by native agarose gel electrophoresis as in Figure S1F. (G) Coomassie-stained SDS-PAGE gel of recombinant proteins used in this study. For Lig4/XRCC4 preps, top band (~100 kDa) is Lig4, bottom band (~50 kDa) is XRCC4, and other bands are minor contaminants.

**Figure S2.**
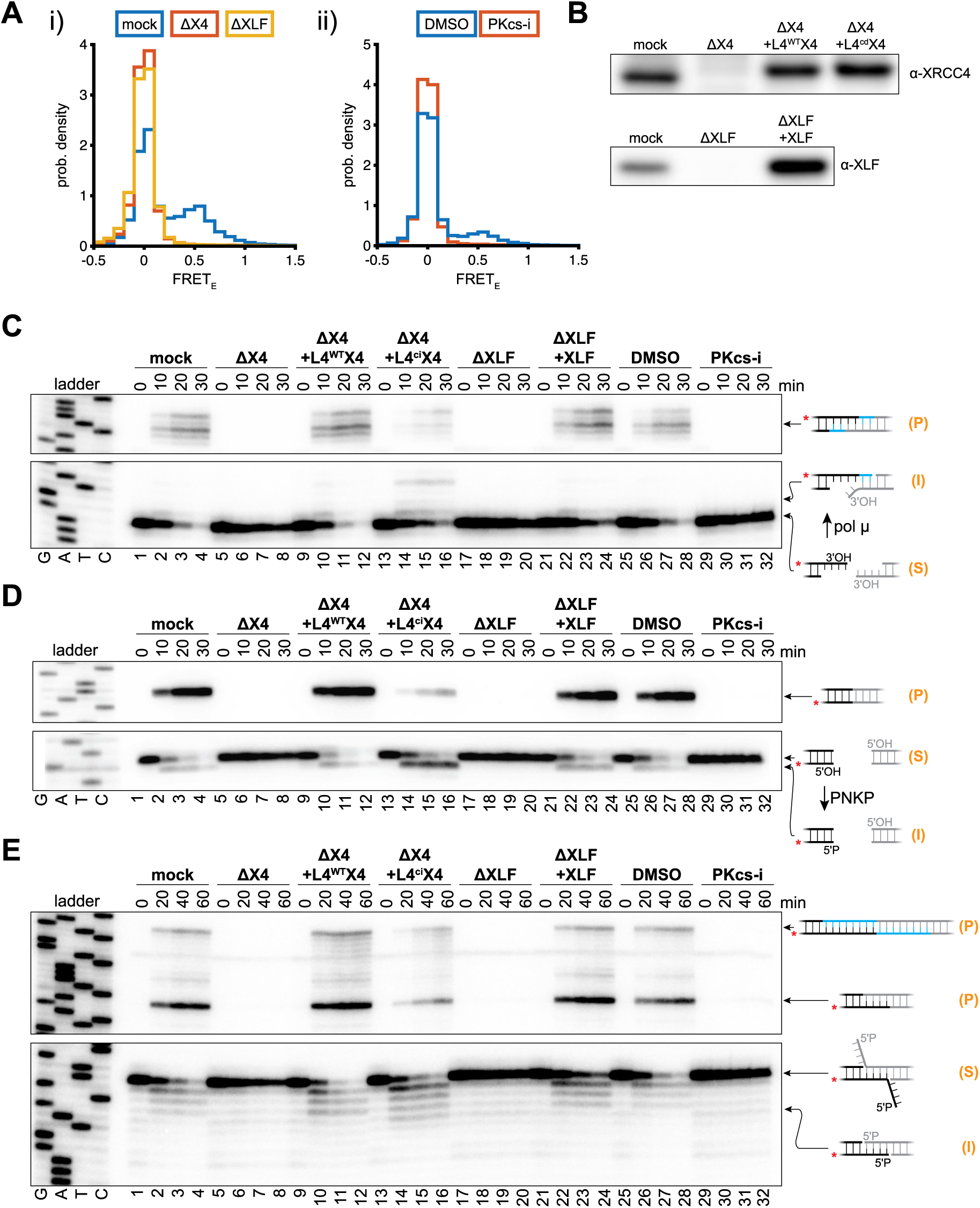
Disruption of short-range synapsis inhibits end processing by pol μ, PNKP, and a 5′→3′ exonuclease. Related to Figure 3. (A) The single-molecule DNA substrate for polymerase activity was immobilized on a glass coverslip in a microfluidic chamber as in Figure 4A, and the indicated extracts were added to the chamber. To minimize photobleaching, a field of view within the chamber was imaged for 30 sec before moving to a new field for a total imaging time of 7.5 min. Panels (i) and (ii) show histograms of apparent FRET efficiency for the indicated extracts. These results demonstrate that DNA-PKcs kinase activity, Lig4-XRCC4, and XLF are required for short range synapsis of these ends. (B) Western blot analysis of XRCC4- and XLF-immunodepleted extracts used here and in Figure 3. Recombinant Lig4-XRCC4 and XLF were added to final concentrations of 50 nM and 100 nM, respectively. (C to E) Denaturing PAGE analysis of end processing and joining of the depicted DNA ends in the indicated extracts, as in Figure 3. S, input substrate; I, processing intermediate; P, ligation product.

**Figure S3.**
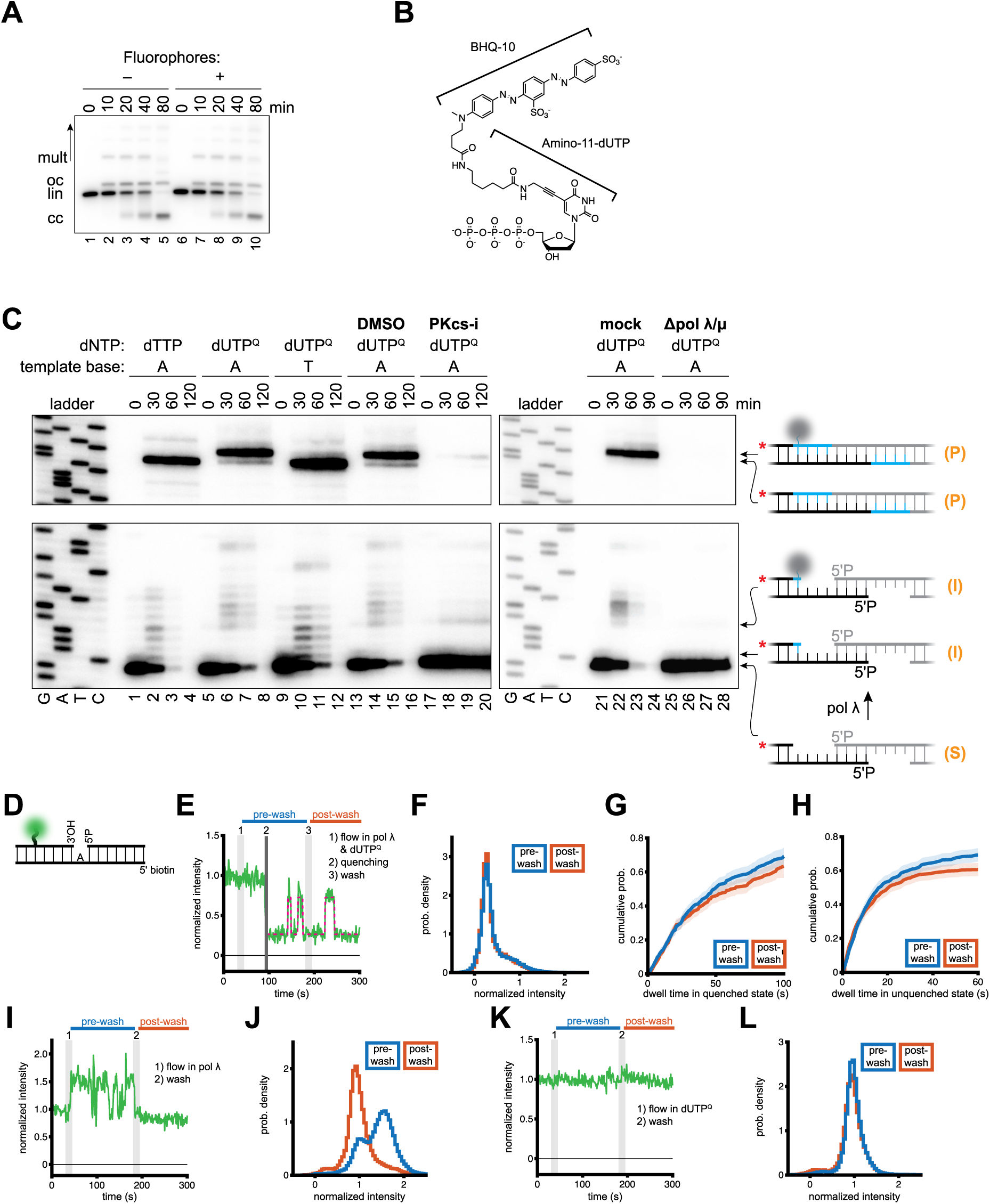
Characterization of the single-molecule polymerase activity assay. Related to Figure 4. (A) Radiolabeled single-molecule DNA substrates with ends as depicted in Figure 4A with or without donor and acceptor fluorophores were added to egg extracts. Reaction samples were stopped at the indicated times and analyzed by native agarose gel electrophoresis and autoradiography. These results indicate that the addition of fluorophores has no significant effect on joining of this substrate. mult, multimers; oc, open circular plasmid; lin, linear; cc, closed circular plasmid. (B) Chemical structure of dUTP^Q^, which was generated by coupling an NHS-ester derivative of BHQ-10 to amino-11-dUTP. (C) Denaturing PAGE analysis of end processing and joining of depicted DNA ends under the indicated conditions. Gray circles in cartoons represent dUTP^Q^, whose incorporation results in slower migration of DNA strands relative to dTTP incorporation. S, input substrate; I, processing intermediate; P, ligation product. (D) Oligonucleotide substrate used to characterize the properties of dUTP^Q^ quenching (see Methods for sequence). This substrate was immobilized to a glass coverslip in a microfluidic chamber as in Figure 4A. Buffered solutions containing recombinant pol λ and/or dUTP^Q^ were introduced to the chamber as indicated below. Green circle, Cy3 donor. The fluorophore is positioned five bases from the gap to match the configuration of the substrate in Figure 4A. (E) Two possible sources of the fluorescence fluctuations observed after the initial quenching event (see Figure 4B for example) are (i) “blinking” of BHQ-10 after incorporation (analogous to fluorophore blinking, but in this case, BHQ-10 would occasionally enter a “dark” state in which it is no longer able to accept photons from Cy3, resulting in increased fluorescence intensity) or (ii) repeated futile dUTP^Q^ incorporation attempts, in which pol λ and dUTP^Q^ bind to the primer-template junction, resulting in quenching, and dissociate without incorporation. To distinguish between these possibilities, we asked whether these fluctuations depended on the presence of pol λ. We began imaging in buffer alone before flowing in buffer containing both 50 nM pol λ and 50 μM dUTP^Q^ (1), which frequently resulted in quenching (2). We then washed the chamber with buffer alone to remove pol λ and dUTP^Q^ (3). As shown in this example, we frequently observed fluorescence fluctuations after removing pol λ and dUTP^Q^, consistent with model (i) and arguing against model (ii). See below for analysis of all molecules. Green, Cy3 intensity, normalized to the average intensity before pol λ/dUTP^Q^ introduction; magenta dotted line, two-state hidden Markov model idealized trajectory (see G and H). (F) Histograms of normalized Cy3 intensity for molecules with a detected quenching event. Blue, the period from initial quenching until the wash step (pre-wash); red, the period from the wash step until the end of observation or censoring due to photobleaching (post-wash). The histograms are superimposable, indicating that removal of pol λ/dUTP^Q^ has no effect on the population of quenched and unquenched states. *n* = 631 molecules with detected quenching event from 2 independent experiments. (G and H) Kinetic analysis of dwell times in the quenched and unquenched states following an initial quenching event. Cy3 intensity following the initial quenching event was fit to a two-state hidden Markov model (magenta dotted line in (E)) to analyze dwell times. Graphs display the cumulative probability distribution of dwell times in pre- and post-wash periods for quenched (G) and unquenched states (H). Pre- and post-wash dwell times are very similar, indicating that removal of pol λ/dUTP^Q^ has minimal effect on the kinetics of fluorescence fluctuations following quenching. (G), *n* = 792 (blue); 700 (red); (H), *n* = 739 (blue); 757 (red). (I) To verify that the wash step effectively removed pol λ, we took advantage of protein-induced fluorescence enhancement (PIFE) observed upon pol λ binding to the oligonucleotide substrate. Upon introducing pol λ (50 nM) without dUTP^Q^ to the chamber, we characteristically observed dynamic increases in Cy3 intensity, and these were eliminated by the wash step. This panel displays a representative example trajectory, with Cy3 intensity normalized to the average intensity of the period before pol λ introduction. (J) Histograms of normalized Cy3 intensity for all molecules. The wash step effectively removed the high-fluorescence population seen in the pre-wash histogram. (K) Example trajectory of normalized Cy3 intensity with dUTP^Q^ (50 μM) alone. (L) Histograms of normalized Cy3 intensity for all molecules demonstrating that introduction of dUTP^Q^ alone had no effect on Cy3 intensity.

**Figure S4.**
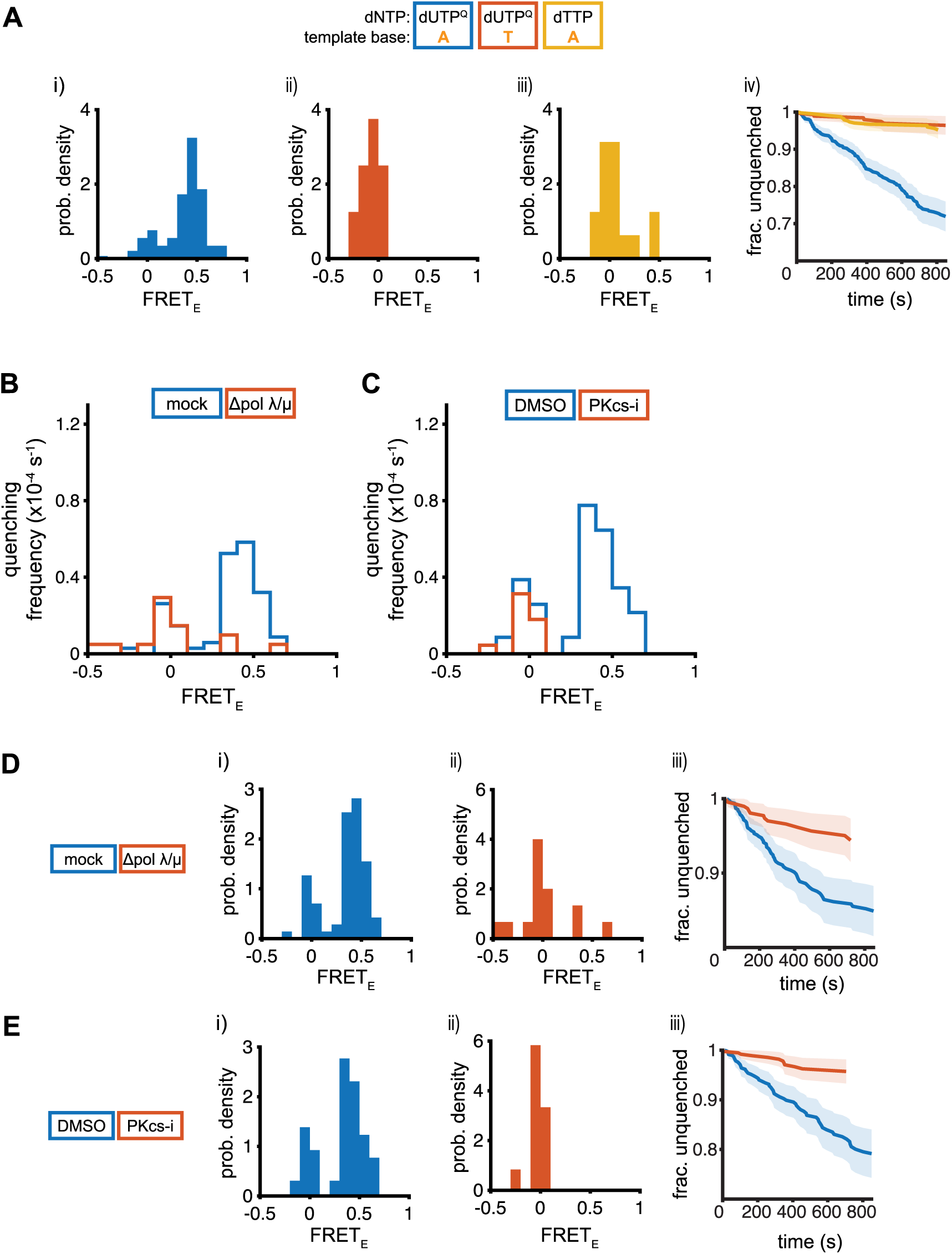
Additional analysis of single-molecule polymerase assay. Related to Figure 4. (A) Extension of Figure 4C and 4D. (Ai to Aiii) Histograms of FRET efficiency immediately prior to identified quenching events under indicated conditions, as in Figure 4C, represented as probability density. (Aiv) survival kinetics (Kaplan-Meier estimate) of un-quenched molecules under the indicated conditions. The x-axis indicates the time from the time extract addition to the identified quenching events. Shaded areas, 95% confidence intervals. *n* = 145 events (blue); 8 events (red); 16 events (yellow). (B) Frequency of quenching events as a function of FRET efficiency immediately prior to quenching, as in Figure 2C, for pol λ/μ-depleted extracts. Blue, mock-depleted extracts (*n* = 71 events from 8 independent experiments); red, pol λ- and pol μ-depleted extracts (*n* = 15 events from 5 independent experiments). See Figure S4D for histograms represented as probability density and kinetic analysis. (C) Frequency of quenching events as a function of FRET efficiency immediately prior to quenching, as in Figure 2C, for DNA-PKcs_i_-treated extracts. Blue, DMSO-treated extracts (*n* = 65 events from 7 independent experiments); red, DNA-PKcs_i_-treated extracts (*n* = 12 events from 6 independent experiments). See Figure S4E for histograms represented as probability density and kinetic analysis. (D and E) Extensions of Figure S4B and S4C, respectively. (Di to Dii; Ei to Eii) Histograms of FRET efficiency immediately prior to identified quenching events under indicated conditions, as in Figure 4C, represented as probability density. (Diii and Eiii) Survival kinetics (Kaplan-Meier estimate) of un-quenched molecules under the indicated conditions. The x-axis indicates the time from the time extract addition to the identified quenching events. Shaded areas, 95% confidence intervals. *n* = 71 events (D, blue); 15 events (D, red); 65 events (E, blue); 12 events (E, red).

**Figure S5.**
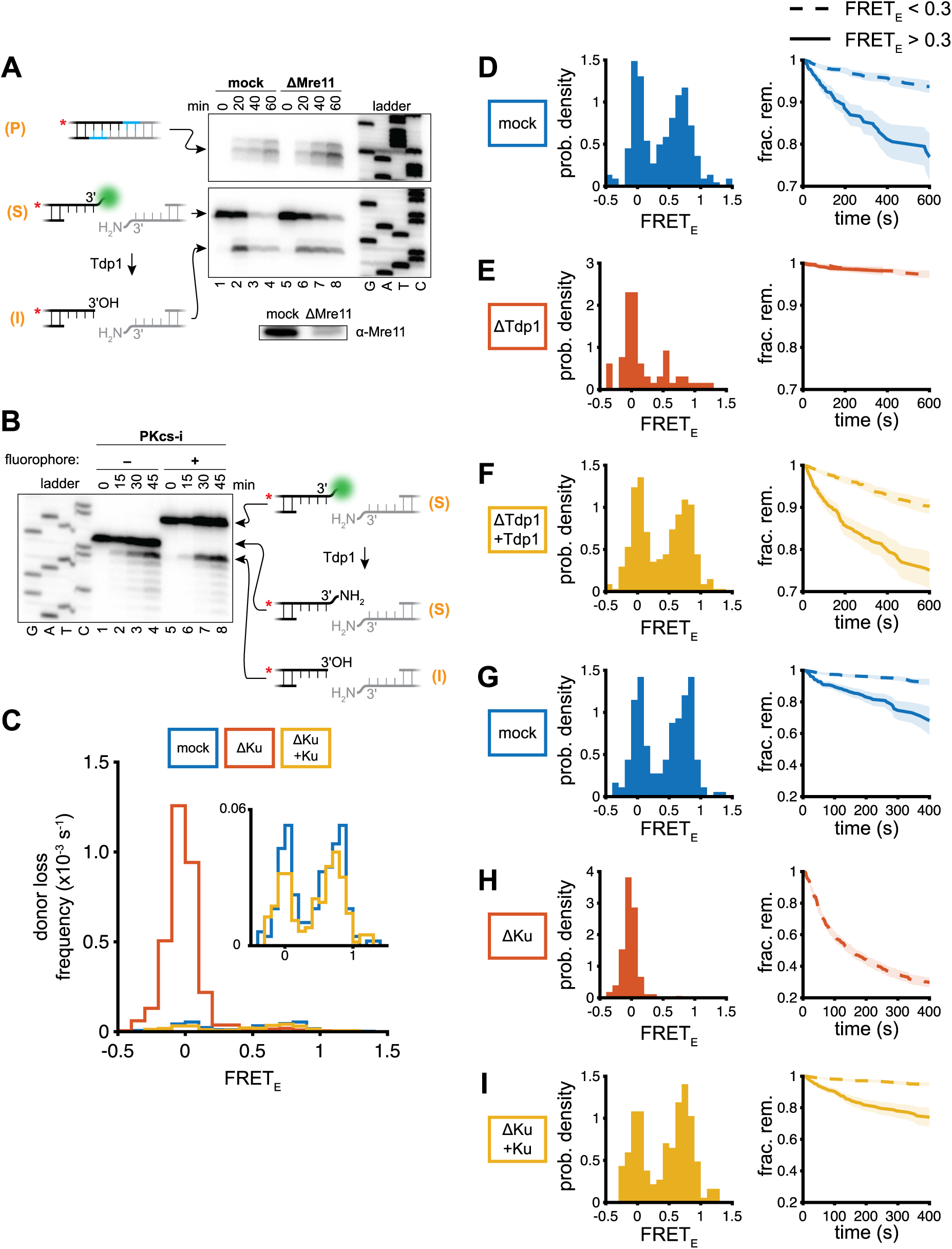
Characterization of the single-molecule assay for Tdp1 activity. Related to Figure 5. (A) Denaturing PAGE analysis, as in Figure 2B, of end processing and joining of the depicted substrate in mock- and Mre11-depleted extracts. Lower panel, Western blot analysis of immunodepleted extracts. We routinely observed ~15% Mre11 remaining following the immunodepletion procedure, but the immunodepletion markedly improved retention of fluorescent DNA molecules on the coverslip surface in single-molecule experiments and stabilized input DNA against resection in ensemble experiments (compare total lane intensity, lanes 4 and 8), indicating Mre11 was functionally depleted. Green circle, Cy3B fluorophore. S, input substrate; I, processing intermediate; P, ligation product. (B) Denaturing PAGE analysis, as in Figure 2B, of end processing and joining of the depicted substrate with and without coupled Cy3B in PKcs-i-treated, Mre11-depleted extracts. Because joining is inhibited by addition of PKcs-i, only unjoined products are shown. S, input substrate; I, processing intermediate; P, ligation product. (C) Frequency of donor loss events as a function of FRET efficiency immediately prior to donor loss events under the indicated conditions. Blue, mock-depleted extracts (*n* = 183 events from 2 independent experiments; 8.3×10^4^ total observations); red, Ku-depleted extracts (*n* = 710 events from 2 independent experiments; 3.7×10^4^ total observations); yellow, Ku-depleted extracts supplemented with 300 nM recombinant Ku (*n* = 185 events from 2 independent experiments; 1.1×10^5^ total observations). See Figure S5G-S5I for kinetic analysis and histograms represented as probability density. (D to F) Extensions of Figure 5C and 5D. Mock immunodepletions in (D) were performed in parallel with Tdp1 immunodepletions in (E) and (F). *n* = 125 events (D, solid); 97 events (D, dashed); 18 events (E, solid); 47 events (E, dashed); 181 events (F, solid); 158 events (F, dashed). (G to I) Extensions of Figures 5E and S5C. Mock immunodepletions in (G) were performed in parallel with Tdp1 immunodepletions in (H) and (I). Left panels, histograms of FRET efficiency immediately prior to identified donor loss events under the indicated conditions represented as probability density. Right panels, donor survival kinetics (Kaplan-Meier estimate) under the indicated conditions. The x-axis indicates dwell time in the high-FRET (solid lines) or low-FRET (dashed lines) state prior to donor loss. Shaded areas, 95% confidence intervals. *n* = 107 events (G, solid); 76 events (G, dashed); 687 events (H, dashed; 23 events were detected with FRET_E_ > 0.3 but are not plotted); 113 events (I, solid); 72 events (I, dashed).

**Figure S6.**
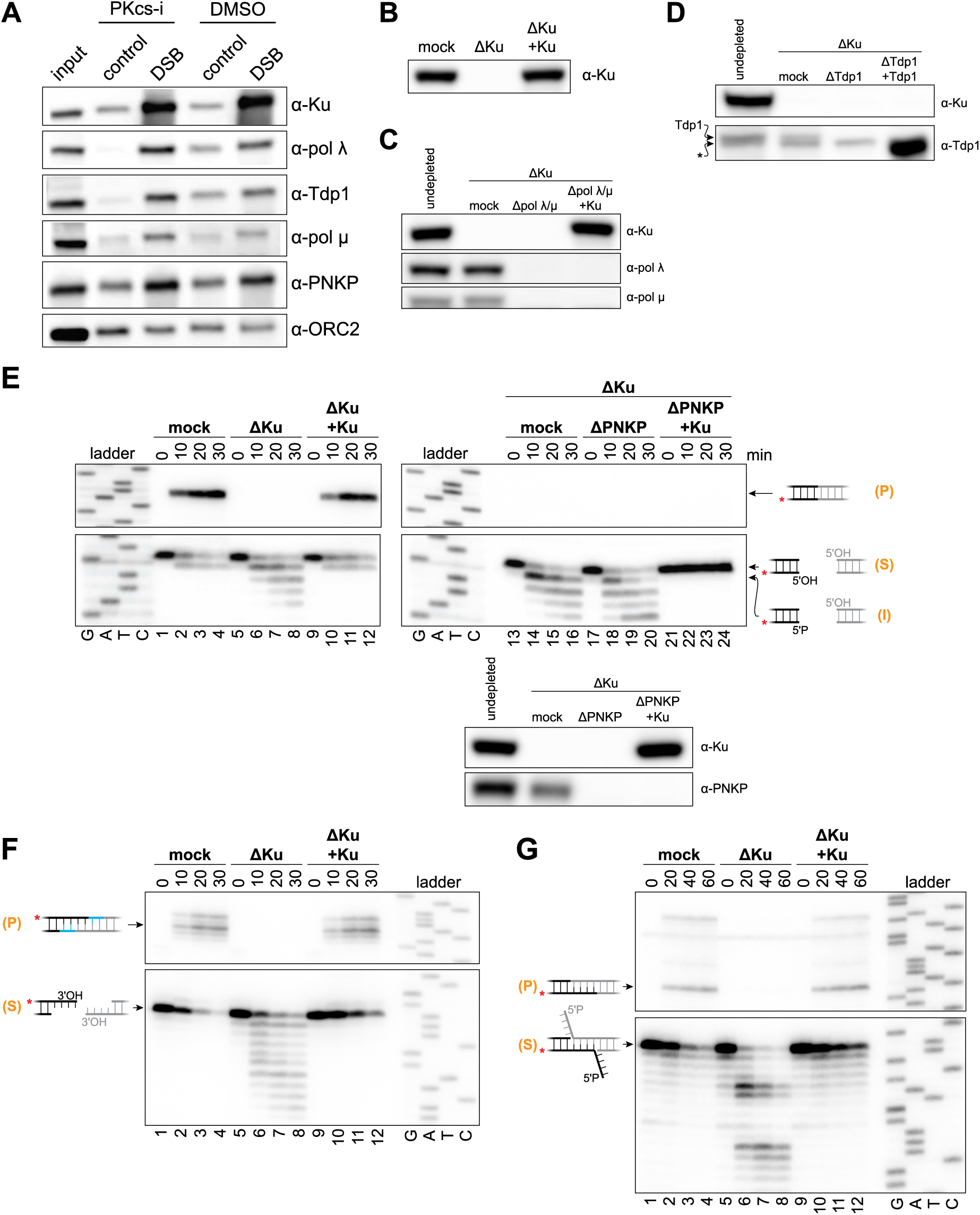
Ku protects DNA ends from premature and off-pathway processing. Related to Figure 6. (A) Representative Western blot analysis used to generate graphs in Figure 6A. Band intensities were measured in ImageJ and normalized to the ORC2 loading control. Fold-enrichment was calculated from the ratio of the normalized band in intensity in the DSB condition relative to that of the intact control. (B to D) Western blot analysis of immunodepleted extracts used here and in Figure 6. Recombinant Ku and Tdp1 were added as indicated to final concentrations of 300 nM and 40 nM, respectively. C, nonspecific band in the Tdp1 immunoblot is marked with an asterisk. (E to G) Denaturing PAGE analysis of end processing and joining of the depicted DNA ends in the indicated extracts, as in Figures 6B and 6C. Lower panels in E, Western blot analysis of immunodepleted extracts. In D, 5′ phosphorylation was observed when both Ku and PNKP were depleted (lanes 17 to 20), but not when recombinant Ku was added back (lanes 21 to 24). A likely explanation is that undetectable residual PNKP is able to phosphorylate unprotected ends when Ku is absent but not when Ku is present. We cannot rule out the possibility that a different enzyme phosphorylates ends in the absence of Ku, but PNKP is the only reported vertebrate enzyme with this biochemical activity. S, input substrate; I, processing intermediate; P, ligation product.

**Figure S7.**
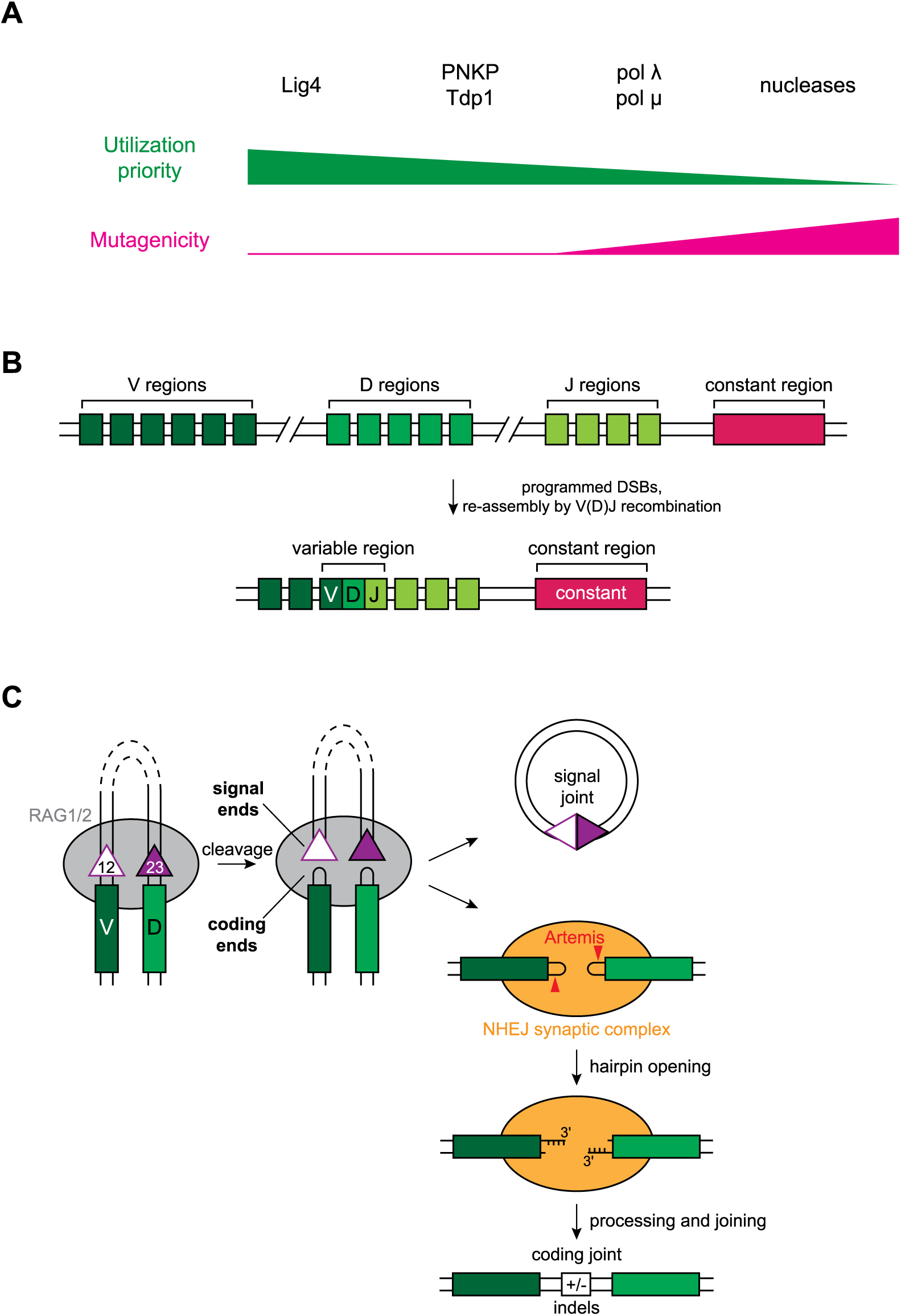
Models of DNA end processing and synapsis. Related to Figure 7. (A) Hierarchical model of DNA end processing during NHEJ of spontaneous DSBs, as described in the main text. (B) Simplified overview of rearrangement of the germline immunoglobulin heavy chain locus. During lymphocyte development, V, D, and J gene segments are rearranged into exons encoding the variable regions of immune receptors. This process involves programmed DSBs that are repaired by NHEJ (Alt et al., 2013; Gellert, 2002). (C) Simplified schematic of V-to-D joining during V(D)J recombination. Proper reorganization is ensured by synapsis of appropriate RSSs according to the “12/23 rule” (Alt et al., 2013). RSSs consist of a highly conserved heptamer sequence, a spacer of either 12 nucleotides (“12RSS”) or 23 nucleotides (“23RSS”), and a conserved nonamer sequence. V and J segments are flanked by 23RSSs, and D segments are flanked by 12 RSSs. In order to cleave DNA, RAG must bind both a 12RSS and a 23RSS, thereby ensuring V-to-D and D-to-J joining (Alt et al., 2013; Gellert, 2002). RAG then introduces DSBs, forming hairpin-capped “coding” ends and blunt “signal” ends that are joined by NHEJ to form V(D)J exons and a circular signal joint product, respectively.

**Table S1.**
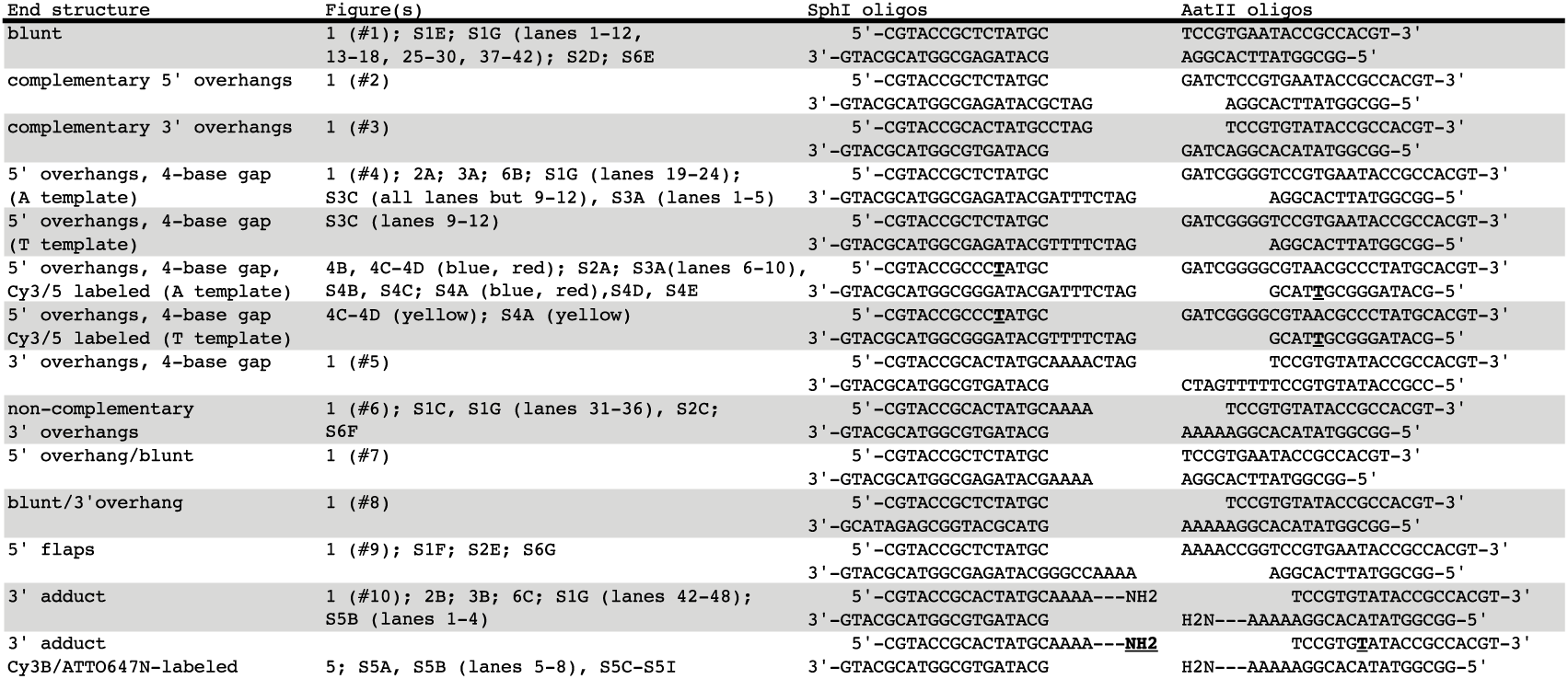
Oligonucleotide duplexes used to generate DNA substrates. DNA substrates were generated as diagrammed in Figure S1A and described in Methods. Bold and underlined characters indicate the position of fluorophore attachment. NHS ester dye derivatives were conjugated via either an internal C6-amino dT (IDT code /iAmMC6T/) or a 3′ amino modifier (IDT code /3AmMO/, structure shown in Figure 1B, #10).

**Data S1. Analysis of aligned sequencing reads.** Sheets labels correspond to DNA ends depicted in Figure 1B.

## Materials and Methods

All ensemble experiments were performed at least twice, with a representative result shown. The sample size and number of individual single molecule experiments can be found in the figure legends. All oligonucleotides were synthesized by Integrated DNA Technologies, Inc. Fluorophores: Sulfo-Cyanine3 NHS ester (Lumiprobe); Sulfo-Cyanine5 NHS ester (Lumiprobe); Cy3B NHS ester (GE Healthcare); ATTO 647N NHS ester (ATTO-TEC). Unless othwerwise noted, DNA-PKi indicates addition of NU7441 (R&D Systems, Inc.) to a final concentration of 50 μM.

### Egg extract preparation

High-speed supernatant (HSS) of egg cytosol was prepared as described (Lebofsky et al., 2009). The female frogs used to produce oocytes were cared for by the Center for Animal Resources and Comparative Medicine at Harvard Medical School (AAALAC accredited). Work performed for this study was in accordance with the rules and regulations set by AAALAC. The Institutional Animal Care and Use Committee (IACUC) of Harvard Medical School approved the work.

### Preparation of DNA substrates

A schematic of DNA substrate preparation is depicted in Figure S1A, and oligonucleotides are listed in Table S1. pBMS6, a derivative of pBlueScript, was linearized with SphI and AatII (New England Biolabs). The resulting 2977 bp fragment was separated on a 1x TBE agarose gel and recovered by electroelution and ethanol precipitation. Duplex oligonucleotide adapters were ligated to each side of this backbone fragment to generate the described DNA substrates.

To generate duplex adapters, oligonucleotide stocks (10 μM) in annealing buffer (10 mM Tris, pH 8.0, 50 mM NaCl, 1 mM EDTA) were combined in equal volumes and annealed by heating to 95°C for 2 min then slowly cooling to room temperature in 1°C steps lasting 60 s. One duplex adapter (oligos 1-2 in Figure S1A) contained an overhang complementary to that generated by SphI; similarly, the second duplex adapter (oligos 3-4) contained an overhang complementary to that generated by AatII. Oligonucleotides (1) and (4) were phosphorylated with T4 PNK (New England Biolabs) to allow ligation to the backbone fragment. Oligonucleotides (2) and (3) were not phosphorylated at the annealing stage.

To generate most radiolabeled DNA substrates, γ-^32^P-ATP (Perkin Elmer) was used to phosphorylate oligonucleotide (1). To label the opposite strand (e.g., Figure 2C), the backbone fragment was treated with Shrimp Alkaline Phosphatase (NEB) prior to phosphorylation with γ-^32^P-ATP. In both cases, ends were first phosphorylated for 30 min with γ-^32^P-ATP, and subsequently 50 μM cold ATP was added and the reaction was allowed to proceed a further 15 min to ensure maximal phosphorylation.

Duplex adapters (250 nM) were ligated to the backbone fragment (25 nM) with T4 DNA ligase. The desired ~3 kb product was separated on a 1x TBE agarose gel and recovered by electroelution and ethanol precipitation (fluorescent DNA substrates) or with the QIAquick gel extraction kit (Qiagen).

Single-molecule DNA substrates were generated in the same way using fluorescently labeled oligos (prepared as described below). Subsequently, the recovered DNA was treated with Nt.BbvCI to introduce two nicks on the same stand near the middle of the molecule, thereby allowing removal of a 25-mer oligonucleotide. A 10-fold molar excess of an internally biotinylated, 5′-phosphorylated oligonucleotide with the same sequence was then added to the digestion mixture, and the mixture was subjected to the same annealing protocol as above, allowing insertion of the biotinylated oligo. The mixture was then treated with T4 DNA ligase, separated on a 1x TBE agarose gel, and the appropriate band was recovered by electroelution and ethanol precipitation.

Finally, all substrates, with the exception of that shown in Figures 2C, S1G (lanes 7-12), S2D, and S6E, were phosphorylated with T4 PNK prior to use.

### Preparation of fluorescently-labeled oligonucleotides

Amino-modified synthetic oligonucleotides (see Table S1) were reacted with NHS-ester fluorophore derivatives. A typical labeling reaction contained 46 μL labeling buffer (100 mM sodium tetraborate, pH 8.5), 2 μL amino-modified oligo (25 mg/mL stock in water), and 2 μL NHS-ester fluorophore (50 mg/mL stock in DMSO). The reaction was allowed to proceed overnight at room temperature in the dark. The mixture was ethanol precipitated to remove excess fluorophore. The pellet was resuspended in ~10 μL Gel Loading Buffer II (Invitrogen) and subjected to denaturing PAGE electrophoresis on a 20% Urea-PAGE gel. The band corresponding to the labeled oligo was excised and crushed by centrifugation through a 1.7 mL microcentrifuge tube with a hole in the bottom made with an 18-gauge needle. 500 μL TE buffer (10 mM Tris, pH 8.0, 0.1 mM EDTA) was added to the crushed gel slice, and the mixture was frozen in liquid nitrogen, rapidly thawed in warm water, and rotated overnight at room temperature in the dark. The solution was collected, and the labeled oligo was recovered by ethanol precipitation.

### Preparation of dUTP^Q^

dUTP^Q^ (Figure S3B) was prepared by conjugating Amino-11-dUTP (Lumiprobe) to BHQ-10 succinimidyl ester (Biosearch Technologies). In a typical reaction, 2 μL 50 mM Amino-11-dUTP and 2.5 μL 50 mM BHQ-10 succinimidyl ester were mixed with 45.5 μL 100 mM sodium tetraborate, pH 8.5, and the reaction was allowed to proceed overnight at room temperature in the dark. The mixture was separated on a HiTrapQ column (GE Healthcare) using an AKTA pure chromatography system (GE Healthcare). The mixture was diluted 10-fold in solvent A (10% acetonitrile in water) and loaded onto the column, followed by a gradient of 0-100% solvent B (3 M LiCl) over 20 column volumes. The elution profile of the mixture showed a single additional peak with absorbance at 516 nm relative to the elution profiles of Amino-11-dUTP and BHQ-10 succinimidyl ester alone. Fractions corresponding to this peak were pooled and added to 30 volumes ice-cold acetone to precipitate dUTP^Q^, which was collected by centrifugation, washed with cold acetone, and resuspended in 10 mM Tris, pH 8.0.

### Protein expression and purification

Ku70/80 and XLF were gifts from T. Graham and were purified as described(Graham et al., 2016). A Coomassie-stained SDS-PAGE gel of recombinantly-purified proteins is shown in Figure S1H.

#### General procedure for purification of H_6_-SUMO-tagged proteins

Cells were grown at 37°C to O.D. 0.5-0.7 in LB broth (typically 2 L) and moved to 18°C for overnight (~16 hr) induction with 1 mM IPTG. Cells were harvested by centrifugation and resuspended in ice-cold Ni-NTA lysis buffer (20 mM HEPES, pH 7.5, 400 mM NaCl, 20 mM imidazole, 10% glycerol, 1 mM DTT; 20 mL per L of culture). Resuspended cells were lysed by sonication in the presence of cOmplete Protease Inhibitor Cocktail, EDTA-free (Roche). Lysates were clarified by centrifugation at 20,000 rcf for 60 min, and the supernatant was added to Ni-NTA agarose resin (0.5 mL bed volume per liter of cell culture; Qiagen) equilibrated in Ni-NTA lysis buffer and agitated by rocking at 4°C for 60 min. Resin was collected by centrifugation, washed once with 10 mL Ni-NTA lysis buffer, resuspended in ~10 mL Ni-NTA lysis buffer and applied to a polypropylene gravity flow column. The resin was washed twice with 10 mL Ni-NTA lysis buffer, and bound proteins were eluted with 1 mL fractions of Ni-NTA elution buffer (Ni-NTA lysis buffer with 250 mM imidazole). Fractions containing protein (determined by Bradford assay; Biorad) were concentrated using a 10,000 MWCO centrifugal filter (Amicon) and desalted using a PD-10 column (GE Healthcare) equilibrated in Ulp1 cleavage buffer (20 mM HEPES, pH 7.5, 150 mM NaCl, 10% glycerol, 1 mM DTT) according to the manufacturer’s instructions. The H6-SUMO tag was removed by treatment with H_6_-tagged Ulp1 protease (~30 μg/mL final concentration) as described for each individual protein below. Imidazole (20 mM final concentration) was added to the reaction mixture, which was then passed over a 0.5 mL Ni-NTA agarose column equilibrated in Ulp1 reaction buffer supplemented with 20 mM imidazole. The flow-though was collected, including a 1 mL wash to collect protein remaining in the void volume, and subjected to further purification steps as indicated below for each protein.

#### Lig4-XRCC4

*X. laevis* Lig4 and XRCC4 coding sequences (Thermo Scientific CloneIDs 6635763 and 6957895) were subcloned into the pETDuet-1 vector (Novagen), and an N-terminal H_6_- SUMO tag was introduced to Lig4. The resulting plasmid (pBMS49 for wild-type Lig4; pBMS50 for the K278R catalytically-inactive mutant) was transformed into Rosetta II (DE3) cells (Novagen), and overexpression and initial purification were performed as described in “General procedure for purification of H_6_-SUMO-tagged proteins,” with Ulp1 cleavage carried out at room temperature for 2 hr. The flowthrough following clean-up of the Ulp1 cleavage mixture was diluted three-fold with dilution buffer (20 mM HEPES, pH 7.5, 10% glycerol, 1 mM DTT) and purified by anion exchange chromatography (MonoQ 5/50GL, GE Healthcare) on an AKTA pure chromatography system (GE Healthcare). Sample was applied to the column, washed with buffer A (20 mM HEPES, pH 7.5, 50 mM NaCl, 10% glycerol, 1 mM DTT), and eluted with a gradient of 0-60% buffer B (20 mM HEPES, pH 7.5, 1 M NaCl, 10% glycerol, 1 mM DTT) over 20 column volumes. Fractions were analyzed by SDS-PAGE, and those containing Lig4-XRCC4 were pooled and concentrated to ~0.5 mL using a 10,000 MWCO centrifugal filter. The sample was applied to a Superdex 200 Increase column (GE Healthcare) equilibrated in Ulp1 cleavage buffer. Fractions were analyzed by SDS-PAGE, pooled, concentrated, flash-frozen in liquid nitrogen, and stored at ­80°C until use.

#### PNKP

The *X. laevis* PNKP coding sequence was amplified from egg cDNA (gift of T. Graham) and subcloned into an N-terminal H_6_-SUMO tag expression vector. The resulting plasmid (pBMS10) was transformed into Rosetta II (DE3) cells, and overexpression and initial purification were performed as described in “General procedure for purification of H_6_-SUMO-tagged proteins,” except Ulp1 cleavage buffer was supplemented with additional NaCl (300 mM total). Ulp1 cleavage was carried out at 30°C for 4 hr, and the mixture was centrifuged at 20,000 rcf for 15 min to remove precipitates. The flowthrough following clean-up of the Ulp1 cleavage mixture was diluted six-fold with dilution buffer (20 mM HEPES, pH 7.5, 10% glycerol, 1 mM DTT) and purified by cation exchange chromatography (MonoS 5/50 GL, GE Healthcare) on an AKTA pure chromatography system (buffer A: 20 mM HEPES, pH 7.5, 50 mM NaCl, 10% glycerol, 1 mM DTT; buffer B: buffer A with 1 M NaCl). Sample was applied to the column, washed with 15% buffer B, and eluted with a gradient of 15-40% buffer B over 20 column volumes. Fractions were analyzed by SDS-PAGE, and those containing PNKP were pooled and concentrated to ~0.5 mL using a 10,000 MWCO centrifugal filter. The sample was applied to a Superdex 200 Increase column (GE Healthcare) equilibrated in Ulp1 cleavage buffer. Fractions were analyzed by SDS-PAGE, pooled, concentrated, flash-frozen in liquid nitrogen, and stored at −80°C until use.

#### pol λ and pol μ

*X. laevis* pol λ and pol μ coding sequences were amplified from egg cDNA and subcloned into an N-terminal H_6_-SUMO tag expression vector. The resulting plasmids (H_6_-SUMO-pol λ, pBMS25; (H_6_-SUMO-pol μ, pBMS14) were transformed into Rosetta II (DE3) cells, and overexpression and initial purification were performed as described in “General procedure for purification of H_6_-SUMO-tagged proteins,” with Ulp1 cleavage carried out at room temperature for 90 min. The flowthrough following clean-up of the Ulp1 cleavage mixture was applied to a Superdex 200 Increase column equilibrated in Ulp1 cleavage buffer. Fractions were analyzed by SDS-PAGE, pooled, concentrated, flash-frozen in liquid nitrogen, and stored at −80°C until use.

#### Tdp1

The *X. laevis* Tdp1 coding sequence (XGC CloneID 6638604; Dharmacon) was subcloned into N-terminal H_6_-SUMO tag expression vector. The resulting plasmid (pBMS60) was transformed into Rosetta II (DE3) cells, and overexpression and initial purification were performed as described in “General procedure for purification of H_6_-SUMO-tagged proteins” with the following modification: Ulp1 was added to the initial Ni-NTA eluate, and the mixture was dialyzed using 12-14 kDa MWCO dialysis tubing (Spectra/Por) against 1 L Ulp1 cleavage buffer overnight at 4°C. The flowthrough following clean-up of the Ulp1 cleavage mixture was diluted three-fold with dilution buffer (20 mM HEPES, pH 7.5, 10% glycerol, 1 mM DTT) and purified by cation exchange chromatography (MonoS 5/50 GL, GE Healthcare) on an AKTA pure chromatography system (GE Healthcare). Sample was applied to the column, washed with buffer A (20 mM HEPES, pH 7.5, 50 mM NaCl, 10% glycerol, 1 mM DTT), and eluted with a gradient of 0-50% buffer B (20 mM HEPES, pH 7.5, 1 M NaCl, 10% glycerol, 1 mM DTT) over 20 column volumes. Fractions were analyzed by SDS-PAGE, and those containing Tdp1 were pooled and concentrated to ~0.5 mL by binding to a small volume of SP sepharose resin (GE Healthcare) and elution with 50% buffer B. The sample was applied to a Superdex 200 Increase column (GE Healthcare) equilibrated in Ulp1 cleavage buffer. Fractions were analyzed by SDS-PAGE, pooled, concentrated with SP sepharose resin as above, flash-frozen in liquid nitrogen, and stored at −80°C until use.

### Antibodies and immunodepletion

Rabbit polyclonal antibodies raised against the following *X. laevis* proteins were previously described: Ku80 (Graham et al., 2016), XLF (Graham et al., 2016), XRCC4 (Graham et al., 2016), and ORC2 (Walter and Newport, 1997). Rabbit polyclonal antibodies were raised against full-length recombinant *X. laevis* PNKP, pol λ, and pol μ (purified as described above) by Pocono Rabbit Farm & Laboratory, Inc. Rabbit polyclonal antibody against *X. laevis* Artemis was raised against the peptide CKLQHVYKRLAMGDNVL by New England Peptide. Rabbit polyclonal antibody against *X. laevis* Tdp1 was raised against the peptide MDRTSASQQSNYGKC by New England Peptide. Rabbit polyclonal antibody against *X. laevis* Mre11 was raised against the peptide CDDEEDFDPFKKSGPSRRGRR by Bethyl Laboratories.

PNKP, pol λ, and pol μ antibodies were affinity purified from rabbit serum by coupling each full-length recombinant protein to AminoLink Coupling Resin (Thermo Scientific) and following manufacturer’s instructions for IgG purification.

Unless noted otherwise below, all immunodepletions were carried out by the same procedure. 3 volumes of 1 mg/mL affinity-purified antibody (10 volumes for Mre11 antibody) was gently rotated with 1 volume Protein A Sepharose beads overnight at 4°C or 1 hour at room temperature. Beads were washed extensively with ELBS (2.5 mM MgCl2, 50 mM KCl, 10 mM HEPES, pH 7.7, 0.25 M sucrose), and five volumes of egg extract containing 7.5 ng/μL nocodazole were immunodepleted by three rounds of gentle rotation with one volume of antibody-bound beads for 60 min at 4°C. Immunodepleted extracts were either used immediately or flash-frozen in liquid nitrogen. For co-immunodepletions, 3 volumes of each 1 mg/mL affinity-purified antibody (10 volumes for Mre11 antibody) was added to 1 volume Protein A Sepharose beads. Mock depletions were performed using IgG purified from rabbit pre-immune serum using Protein A Sepharose beads.

λ protein phosphatase treatment of endogenous Tdp1 (Figure S1D(ii)) was performed in the following manner. 20 μL extract was subjected to a single round of Tdp1 immunodepletion as described above. Beads were washed twice with ELBS and suspended in 20 μL PMP buffer supplemented with 1 mM MnCl2 (NEB). 200 units λ protein phosphatase (NEB) was added to 10 μL of the suspension, and the reaction was incubated at 30°C for 30 min before addition of one volume 2x reducing Laemmli sample buffer and immunoblotting as described below.

### Immunoblotting

Samples were resolved on Mini-PROTEAN precast gels (Bio-Rad) and transferred to PVDF membranes (Perkin Elmer). Membranes were blocked in 5% nonfat milk in 1x PBST for 60 min at room temperature, then incubated with antibody diluted 1:2000 in 1x PBST containing 1% BSA for 60 min at room temperature or overnight at 4°C. After extensive washing in 1x PBST at room temperature, the membranes were incubated with goat anti-rabbit horseradish peroxidase-conjugated antibody (Jackson ImmunoResearch) diluted 1:20,000 in 5% nonfat milk in 1x PBST for 45 min at room temperature. Membranes were washed extensively in 1x PBST, incubated for ~60 s with HyGLO chemiluminescent HRP antibody detection reagent (Denville), and imaged using an Amersham Imager 600 (GE Healthcare).

### Ensemble NHEJ assays

Ensemble NHEJ assays were conducted at room temperature. Egg extracts were supplemented with the following (final concentration indicated in parentheses): nocodazole (7.5 ng/μL) if nocodazole had not been added during prior immunodepletion; pBMS6 (10 ng/μL); ATP (3 mM); phosphocreatine (15 mM); and creatine phosphokinase (0.01 mg/mL; Sigma). Joining reactions were initiated by addition of 5 ng/μL radiolabeled linear DNA substrate (final concentration, prepared as described above).

For analysis by agarose gel electrophoresis, samples were withdrawn at the indicated times and mixed with a 2.5 volumes agarose stop solution (80 mM Tris, pH 8.0, 8 mM EDTA, 0.13% phosphoric acid, 10% Ficoll, 5% SDS, 0.2% bromophenol blue). Samples were treated with Proteinase K (1.4 mg/mL final concentration) for 60 min at 37°C or room temperature overnight, and products were separated by electrophoresis on a 1x TBE 0.8% agarose gel. Gels were dried under vacuum on a HyBondXL nylon membrane (GE Healthcare) and exposed to a storage phosphor screen, which was imaged with a Typhoon FLA 7000 imager (GE Healthcare).

For analysis by denaturing urea-PAGE, samples were withdrawn at the indicated times and mixed with 10 volumes extraction stop solution (100 mM Tris, pH 8.0, 25 mM EDTA, 0.5% SDS). Samples were treated with RNase A (0.2 mg/mL final concentration) for 30 min at 37°C and then treated with Proteinase K (0.67 mg/mL final concentration) for 60 min at 37°C or room temperature overnight. Samples were phenol-chloroform extracted, ethanol precipitated, resuspended, and simultaneously digested with SacI and KpnI (NEB). Digestion was stopped by addition of 1 volume Gel Loading Buffer II (Invitrogen). A denaturing sequencing gel (6% acrylamide/bis-acrylamide [5% crosslinker], 7 M urea, 40% formamide) was pre-run on a LABREPCO Model S2 apparatus with 0.8x glycerol tolerant gel buffer (20x stock: 1.78M Tris, 0.57 M taurine, 0.01 M EDTA) for 60 min before loading samples, which were heated to 95°C for 2 min and rapidly cooled on ice just prior to loading. Following electrophoresis, the gel was fixed in 20% methanol/10% acetic acid for ~45 min, transferred to Whatman 3 MM Chr filter paper, dried under vacuum, and exposed to a storage phosphor screen, which was imaged with a Typhoon FLA 7000 imager. Sequencing ladders were generated using the Thermo Sequenase Cycle Sequencing kit (Affymetrix) according to the manufacturer’s instructions.

### Deep sequencing of NHEJ products

Joining reactions were carried out as described above for 90 min. Products were separated on a 1x TBE, 0.8% agarose gel and stained with SYBR Gold (Invitrogen). The band corresponding to closed-circular plasmid was excised and extracted with the QIAquick gel extraction kit (Qiagen). A second independent replicate was performed, and the recovered DNA was pooled. This material served as the template for amplification by PCR prior to next-generation sequencing. For each sample, a ~200 bp amplicon was generated from 1×10^6^ template molecules by PCR in 22 cycles using Phusion polymerase (NEB), purified with the QIAquick gel extraction kit, and submitted to Genewiz for the Amplicon-EZ service (paired-end Illumina sequencing) and analysis. Reads were aligned using a 40 bp target sequence centered at the break site and compared to the reference sequence corresponding to direct joining. Two control templates were amplified in parallel: (1) a template of known sequence, to control for errors in amplification and library preparation, and (2) a mixture of three templates in a defined ratio of 48 (“WT”): 1 (1 bp substitution): 1 (12 bp deletion), to ensure relatively rare sequences were recovered. Control (1) returned 98.5% of reads with the expected sequence. Control (2) returned the expected reads in a ratio of 43:1:2. Deviation from the expected 48:1:1 ratio is likely due to slight amplification bias towards the shorter amplicon and heterogeneity in the template, which in this case was synthetic double-stranded DNA fragments. The analysis results for all samples are reported in Data S1.

### Single-molecule microscope and chamber preparation

Samples were imaged with a through-objective TIRF microscope built around an Olympus IX-71 inverted microscope body. 532-nm, and 641-nm laser beams (Coherent Sapphire 532, and Cube 641, respectively) were expanded, combined with dichroic mirrors, expanded again, and focused on the rear focal plane of an oil-immersion objective (Olympus UPlanSApo, 100×; NA, 1.40). The focusing lens was placed on a vertical translation stage to permit manual adjustment of the TIRF angle. Emission light was separated from excitation light with a multipass dichroic mirror, and laser lines were further attenuated with a StopLine 488/532/635 notch filter (Semrock). A home-built beamsplitter (Graham et al., 2017) was used to separate Cy3 emission from Cy5 emission; these two channels were imaged on separate halves of an electron-multiplying charge-coupled device camera (Hamamatsu, ImageEM 9100-13), which was operated at maximum EM gain. An automated microstage (Mad City Labs) was used to position the sample and move between fields of view.

Microfluidic chambers were constructed in the following manner: a Dremel tool with a diamond-tipped rotary bit was used to drill two holes 10 mm apart in a glass microscope slide; PE20 tubing was inserted into one hole and PE60 tubing into the other (Intramedic), and the tubing was cut flush on one side of the slide and fixed in place with epoxy (Devcon) on the other; double-sided SecureSeal Adhesive Sheet (Grace Bio-Labs), into which a 1.5 × 12 mm channel had been cut, was placed on the non-tubing side of the slide, aligning the channel with the holes in the slide. A glass coverslip, functionalized with a mixture of methoxypolyethylene glycol (mPEG) and biotin-mPEG (Laysan Bio, Inc.) as previously described(Graham et al., 2017), was then placed on the second side of the adhesive sheet, and the edges of the coverslip were sealed with epoxy.

Solutions were drawn into the chamber by attaching a 1 mL syringe to the PE60 tubing. Flow cells were incubated with 1 mg/mL streptavidin (Sigma) in PBS for ~2 min. Unbound streptavidin was washed out with ELB150 (2.5 mM MgCl2, 150 mM KCl, 10 mM HEPES, pH 7.7) and biotinylated DNA substrates were incubated in the channel at a concentration yielding appropriate surface density (typically ~ 1 nM, diluted in ELB150). Unbound DNA was washed out with ELB150 and experiments were performed as indicated below.

### Analysis of single-molecule data

Analysis of single-molecule experiments in egg extracts was performed essentially as described (Graham et al., 2016, 2017) using custom MATLAB scripts, with modifications as noted below. Fluorescence intensities were corrected for donor bleedthrough in the acceptor channel, direct acceptor excitation by the 532 nm laser, and differences in donor/acceptor quantum yield and detection efficiency as described (Graham et al., 2016).

### Single-molecule assay for polymerase activity

To remove endogenous dNTPs, 30 μL egg extract was applied to and eluted from a Micro Bio-Spin 6 column (Bio-Rad) equilibrated in ELB150. The buffer-exchanged extract was supplemented with the following (final concentrations in parentheses): nocodazole (7.5 ng/μL) if nocodazole had not been added during prior immunodepletion; intact pBMS6 (10 ng/μL); SphI/AatII-digested pBMS6 (2.5 ng/μL); ATP (3 mM); phosphocreatine (15 mM); creatine phosphokinase (0.01 mg/mL; Sigma); dATP, dCTP, dGTP (50 μM each); dUTPQ or dTTP, as indicated (50 μM); protocatechuic acid (PCA; 5 mM; Sigma); protocatechuate 3,4-dioxygenase (PCD; 0.1 μM; Sigma); and Trolox (6-Hydroxy-2,5,7,8-tetramethylchromane-2-carboxylic acid; 1 mM; Sigma). PCA and PCD constitute an oxygen-scavenging system and Trolox functions as a triplet-state quencher to improve fluorophore performance.

The biotinylated, fluorescent DNA substrate, generated as described above using oligonucleotides listed in Table S1, was immobilized on a glass coverslip in a microfluidic chamber. Extract was introduced to the chamber and images were taken continuously at a rate of 1 frame/s for 900 s, alternating between two frames of 532 nm excitation and one frame of 641 nm excitation. Surface laser power density was measured using a Coherent FieldMate power meter with an OP-2 VIS detector (532 nm: 2 mW/cm^2^; 641 nm: 0.6 mW/cm^2^).

In the experiments shown in Figures 4 and S4, peak selection was performed with a slightly modified protocol: 2D Gaussian fitting was performed on images from both channels, as well as the same images rotated 45 degrees, and spots were accepted for analysis only if the ellipticity of all fits (ratio of major to minor axis width) were below a specified cutoff. The same images were rotated 45 degrees and subjected to the same ellipticity analysis to exclude spots distorted in a diagonal direction. Additionally, peaks with donor and acceptor centroids displaced by more than 1.25 pixels between channels were rejected. Custom MATLAB scripts were used to automatically identify donor and acceptor photobleaching, FRET transitions, and donor quenching. Trajectories were truncated due to donor photobleaching when, during donor excitation, donor and acceptor intensity fell and remained below a threshold set at 25% of initial donor intensity.

Trajectories were truncated due to acceptor photobleaching when, during acceptor excitation, acceptor intensity fell and remained below a threshold set at 25% of initial acceptor intensity. FRET transitions were detected by locating peaks above a set threshold in the first derivative (MATLAB gradient function) of the smoothed FRET signal (MATLAB smooth function). Trajectories of non-photobleached molecules were truncated due to donor quenching when the average donor intensity in a 6-frame window fell below a threshold set at 40% of the average intensity of the preceding 3 frames without a concomitant FRET transition. Quenching frequency in Figure 4 was calculated by dividing the number of quenching events by the total observation time prior to censoring by quenching, donor or acceptor photobleaching, or end of experiment. Observation times were identical for all experiments (900 s). Survival plots in Figure S4 were generated in MATLAB using (1-ecdf) accounting for right-censoring due to donor or acceptor photobleaching, substrate detachment, or end of experiment. Confidence bounds were calculated using Greenwood’s formula. Survival plots in Figure 4E were similarly constructed, but also accounted for right-censoring upon a FRET transition.

For the single-color experiments shown in Figure S3D-S3L, the DNA substrate was generated by annealing the following oligos (with IDT modification codes) as described above:

CGTACCGCCC/iAmMC6T/ATGC (Cy3-labeled)

/5phos/GCAGGACGAAGCCGA

/5Biosg/TCGGCTTCGTCCTGCAGCATAGGGCGGTACG.

Peaks were selected and background-corrected intensities extracted using the iSMS MATLAB software package (Preus et al., 2015). A custom MATLAB script was used to truncate trajectories upon photobleaching and automatically identify quenching events, detected using a threshold set at 40% of the average intensity prior to the first flow-in step (see Figure S3E, S3I, and S3K). For Figure S3E, S3G, and S3H, intensities were extracted for molecules with a detected quenching event from the quenching event to the end of the trajectory, either due to photobleaching or experiment termination. These intensities were globally fitted to a two-state Hidden Markov Model using the ebFRET MATLAB software package (van de Meent et al., 2014) with default parameters. Cumulative probability plots in Figure S3G and S3H were generated from this fit using the MATLAB ecdf function with right-censoring and confidence bounds with Greenwood’s formula.

### Single-molecule assay for Tdp1 activity

For the experiments shown in Figures 5 and S5, extracts were depleted of Mre11 (and Tdp1 or Ku, as indicated) as described above. Extracts were supplemented with the following (final concentrations in parentheses): pBMS6 (10 ng/μL); un-labeled, linear DNA substrate generated as described above using oligonucleotides listed in Table S1 (2.5 ng/μL); ATP (3 mM); phosphocreatine (15 mM); creatine phosphokinase (0.01 mg/mL); PCA (5 mM); PCD (0.1 μM); and Trolox (1 mM).

The biotinylated, fluorescent DNA substrate, generated as described above using oligonucleotides listed in Table S1, was immobilized on a glass coverslip in a microfluidic chamber. Extract was introduced to the chamber and 100 ms exposures were taken stroboscopically every 1 s. The stage was moved after every exposure in a square pattern, allowing imaging of 4 fields of view in one experiment. 532 nm (surface power density 2 mW/cm^2^) and 641 nm excitation (surface power density 1.5 mW/cm^2^) were alternated such that for each field of view two frames of 532 nm excitation and one frame of 641 nm excitation were recorded with each frame separated by 4 s.

Peak selection was slightly modified to reject peaks whose donor and acceptor centroids were displaced by more than 1 pixel and peaks whose initial intensity fell outside a set window characteristic of single molecules. Custom MATLAB scripts were used to automatically identify acceptor photobleaching, FRET transitions, and donor loss/photobleaching. Trajectories were truncated due to acceptor photobleaching when, during acceptor excitation, acceptor intensity fell and remained below a threshold set at 25% of initial acceptor intensity. FRET transitions were detected by locating peaks above a set threshold in the gradient of the smoothed FRET signal. Trajectories were truncated due to donor loss/photobleaching when, during acceptor excitation, the sum of donor and acceptor intensity fell and remained below a threshold set at 25% of initial donor intensity.

Events in which donor and acceptor intensity fell below their respective thresholds simultaneously, likely due to detachment of the DNA molecule from the coverslip, were not scored as donor loss events. Donor loss frequency in Figure 5 was calculated by dividing the number of donor loss events by the total observation time prior to censoring by donor loss, acceptor photobleaching, or end of experiment. Observation times were identical for all experiments (1800 s). Survival plots in Figures 5E and S5 were generated in MATLAB using (1-ecdf) accounting for right-censoring due to acceptor photobleaching, substrate detachment, end of experiment, or transition to the other FRET state. Confidence bounds were calculated using Greenwood’s formula.

### DNA-pulldown and analysis of protein enrichment on DSBs

#### Substrate generation

A 1 kb long DNA fragment with biotin molecules attached to the 5′ termini (dibiotin-DNA) was generated by PCR amplification using Q5 High-Fidelity Polymerase (NEB), with pAM075 as a template and the 5′-biotinylated primer pair oAM091/oAM092. Biotinylated DNA fragments were isolated by electroelution from a 1.0% 1x TBE-agarose gel, ethanol precipitated, and stored in TE buffer at −20°C.

#### DNA-bead generation

For each biological replicate, 40 µL of streptavidin-coated magnetic beads (Sigma, Dynabeads M-280) were used. Beads were prepared with two washes of 180 µL of 2x binding and wash buffer (2xBWB; 10 mM Tris, pH 7.4, 2 M NaCl, and 20 mM EDTA). In between washes, the beads were centrifuged for 30 seconds at 5000 rpm in a microfuge (Eppendorf 5418) and then placed on a magnetic rack. The magnetic beads were resuspended in 180 µL of 1xBWB containing 30 nM dibiotin-DNA and incubated for 20 minutes on a rotisserie at 25°C. DNA-bound beads were washed twice with 180 µL of 2xBWB, twice with 80 µL of 1x Cutsmart buffer (NEB), and then resuspended in 80 µL of 1x Cutsmart buffer. The DNA-bead suspension was then split in half to prepare plasmid and DSB samples. 1 µL of Xmn I restriction enzyme (NEB) was added to the DSB sample and then both samples were incubated at 37°C for six hours. Xmn I was removed by three washes of 80 uL 2xBWB. To reduce non-specific binding, the beads were washed with 80 uL of egg lysis blocking buffer (ELB-block; 10 mM HEPES, pH 7.7, 50 mM KCl, 2.5 mM MgCl_2_, 250 mM sucrose, 0.25 mg/mL BSA, 0.03% casein, and 0.02% Tween20). Then, DNA-beads were blocked in 80 µL of ELB-block for 20 minutes on ice and resuspended in 50 µL ELB-block.

#### Pulldown of NHEJ Factors

For each biological replicate, 125 µL of HSS was freshly thawed in the presence of nocadazole at a final concentration of 7.5 ng/µL. To reduce non-specific binding of aggregates to the DNA beads, the HSS was centrifuged (14000 rpm, 10 min., 4°C) in a fixed angle microfuge (Eppendorf 5418). The HSS was then split into two 50 µL samples and the DNA-PKcs inhibitor NU-7441 (R&D Systems, Inc.), stored as 4 mM stock in DMSO, was added to a final concentration of 100 µM for the PKcs-i sample, while an equivalent volume of DMSO was added to the other sample. After a 5 minute incubation on ice, an ATP regeneration system (ARS) was added to ensure constant ATP levels. 30x ARS stock contained 65 mM ATP (Sigma), 650 mM phosphocreatine (Sigma), and 160 ng/mL creatine phosphokinase (Sigma) and was used at 1x final concentration. Circular pBluescript II was added to the DNA-bead samples to a final concentration of 30 ng/µL. NHEJ reactions were initiated by mixing 25 µL of the DNA-bead sample with an equal volume of HSS and incubating for 15 minutes on a rotisserie at 25°C. To isolate DNA-beads and associated proteins, 48 µL of the total reaction was layered over 180 µL of ELB-cushion solution (10 mM HEPES, pH 7.7, 50 mM KCl, 2.5 mM MgCl_2_, 500 mM sucrose) in a 5 × 44 mm micro centrifuge tube (Beckman Coulter) and sedimented by centrifugation (12000 rpm, 45 sec., 4°C) in a horizontal rotor (Kompspin, Ku Prima-18R). The pelleted DNA-beads was then washed with 180 µL of ELB-block and resuspended in 50 µL of 1x reducing Laemmli sample buffer. Input samples for Western blotting were prepared by diluting HSS 1:40 in 1x reducing Laemmli sample buffer.

#### Western Blotting

Samples were separated on a 4-15% precast SDS-PAGE gel (BioRad) for 2 hours at 100 V and transferred to a PVDF membrane for 50 minutes at 85 V at 4°C. Membranes were blocked with 5% powdered nonfat milk dissolved in 1x PBST for 30 minutes and then incubated with primary antibody diluted in 1x PBST containing 2.5% BSA (OmniPur) for 12-16 h at 4°C. Primary antibodies were used at the following concentrations: α-Ku80 1:10000, α-Pol λ 1:2000, α-Pol µ 1:2000, α-PNKP 1:2500, α-TDP1 1:4000, and α-Orc2 1:10000. After extensive washing with 1x PBST, membranes were incubated with goat anti-rabbit-HRP (Jackson ImmunoResearch) secondary antibody diluted 1:10,000 or 1:20,000 in 5 % powered nonfat milk and 1x PBST for 1 hour at room temperature. Membranes were washed again with 1x PBST, incubated with HyGLO Quick Spray (Denville Scientific), and imaged on an Amersham Imager 600 (GE Healthcare).

#### Western blot Quantification

Protein levels were quantified using NIH ImageJ 1.51a software using Orc-2 levels as a reference. All pulldowns were performed as three independent experiments. Quantitative data are expressed as mean ± standard deviation (SD). Statistical analyses were performed using an unpaired Student’s t-test in Windows Excel 2013. P-values ≤ 0.05 were considered significant for all analyses.

## Acknowledgments

We thank R. Scully, T. Graham, I. Rivera, V. Kumar, and members of the Walter and Loparo laboratories for helpful discussions and comments on the manuscript. This work was supported by National Institutes of Health grant R01GM115487 (to J.J.L.) and The Howard Hughes Medical Institute (to J.C.W.). B.M.S. is supported by a postdoctoral fellowship from the Damon Runyon Cancer Research Foundation. A.T.M. is supported by a postdoctoral fellowship from the American Cancer Society. J.C.W. is an investigator of the Howard Hughes Medical Institute.

## Author contributions

B.M.S. performed all experiments, with the exception of DNA pull-down experiments, which were performed by A.T.M. J.J.L., J.C.W., and B.M.S. conceived experiments, analyzed the data, and wrote the paper with input from A.T.M.

